# Congenital Zika Syndrome is associated with interferon alfa receptor 1

**DOI:** 10.1101/715862

**Authors:** Tamiris Azamor, Daniela Prado Cunha, Andréa Marques Vieira da Silva, Ohanna Cavalcanti de Lima Bezerra, Marcelo Ribeiro-Alves, Thyago Leal Calvo, Fernanda de Souza Gomes Kehdy, Fernanda Saloum de Neves Manta, Thyago Gomes Pinto, Laís Pereira Ferreira, Elyzabeth Avvad Portari, Letícia da Cunha Guida, Leonardo Gomes, Maria Elisabeth Lopes Moreira, Elizeu de Carvalho, Cynthia Chester Cardoso, Marcelo Muller, Ana Paula Dinis Ano Bom, Patrícia Cristina da Costa Neves, Zilton Vasconcelos, Milton Ozorio Moraes

**Affiliations:** Laboratório de Hanseníase. Instituto Oswaldo Cruz. Fiocruz, Brazil; Vice-Diretoria de Desenvolvimento Tecnológico. Instituto de Tecnologia em Imunobiológicos, Fiocruz, Brazil; Unidade de Pesquisa Clínica, Instituto Nacional de Saúde da Mulher, da Criança e do Adolescente Fernandes Figueira. Fiocruz, Brazil; Laboratório de Pesquisa Clínica em DST/AIDS. Instituto Nacional de Infectologia. Fiocruz, Brazil; Laboratório de Diagnóstico por DNA. Universidade do Estado do Rio de Janeiro, Brazil; Laboratório de Virologia Molecular. Universidade Federal do Rio de Janeiro, Brazil

**Keywords:** Congenital Zika Syndrome, rs2257167, placenta, type I interferon, type III interferon

## Abstract

**Background:** Host factors that influence Congenital Zika Syndrome (CZS) outcome remain elusive. Interferons have been reported as the main antiviral factor in Zika and other flavivirus infections.

**Methods:** We accessed samples from Zika pregnancies, conducted a case-control study to verify whether interferon alfa receptor 1 (*IFNAR1*) and interferon lambda 2 and 4 (*IFNL2/4*) single nucleotide polymorphisms (SNPs) contribute to CZS newborn outcome and we characterized placenta gene expression profile at term.

**Findings:** Newborns carrying CG/CC genotypes of rs2257167 in *IFNAR1* presented higher risk of developing CZS (OR=3.73; IC=1.36-10.21; *Pcorrected*=0.02646). No association between *IFNL* SNPs and CZS was observed. Placenta from CZS cases displayed lower levels of *IFNL2* and *ISG15* along with higher *IFIT5.* The rs2257167 CG/CC placentas also demonstrated high levels of *IFIT5* and inflammation-related genes.

**Interpretation:** We found CZS to be related with exacerbated type I IFN and insufficient type III IFN in placenta at term, forming an unbalanced response modulated by the *IFNAR1* rs2257167 genotype. These findings shed light on the host-pathogen interaction focusing on the genetically regulated type I / type III IFN axis that could lead to better management of Zika and other TORCH (Toxoplasma, Others, Rubella, Cytomegalovirus, Herpes) congenital infections.

**Funding:** This work was supported by the Instituto Oswaldo Cruz (Rio de Janeiro, Brazil) and by the Instituto de Tecnologia em Imunobiológicos (Rio de Janeiro, Brazil).

**Research in context:** *Evidence before this study:* Levels of type I and type III interferons are genetically controlled and decisively regulate outcome of spontaneous viral infections or response to antiviral treatment. Hepatitis C virus, Yellow Fever and Zika virus belong to the Flaviviridae family and elicit similar host immune responses. Congenital Zika Syndrome presents well-known risk factors, mainly the first trimester of pregnancy as well as social and nutritional factors, however, these do not entirely explain abnormal outcomes.

*Added value of this study:* We conducted a case-control study to evaluate SNPs in type I and III interferon genes using samples from newborns and mothers who had zika infection during pregnancy. We have shown that newborn interferon type I background contributes to the development of abnormal CSZ. This specific genetic makeup regulates placental immunological responses and prevents an exacerbated type I, and lack of type III, interferon response in syndromic cases.

*Implications of all the available evidence:* Our study suggests an important factor regulating the host-pathogen interaction during Zika virus (ZIKV) infections in humans. During pregnancy, genetic variations play a role in balancing tissue-specific type I and III interferons during ZIKV congenital infection influencing fetal neurological damage. Custom pharmacological interventions could be used to modulate immunity and inflammation towards protective responses.

**Graphical abstract:** 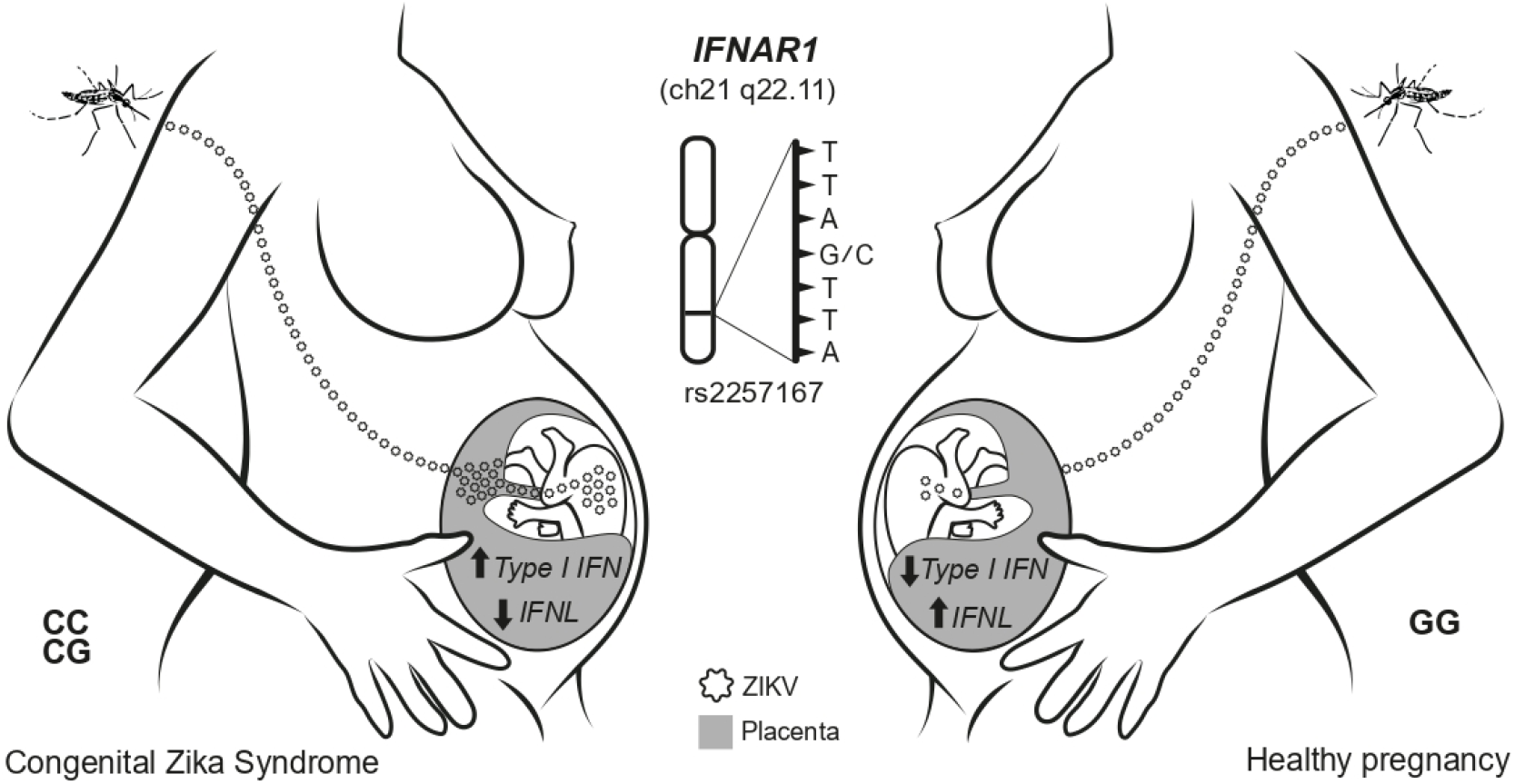

## 1. Introduction

Zika virus (ZIKV) is a single stranded positive-sense RNA virus that belongs to the Flaviviridae family. Zika infection is mostly asymptomatic or associated with mild symptoms. After the outbreak in the Americas in 2015, the virus spread across 59 countries and more than 500.000 suspected cases were reported [1,2]. In a short while, there was a rise in cases of congenital abnormalities, including cerebral anomalies, congenital contractures, ocular alterations among other neurological abnormalities known as Congenital Zika Syndrome (CZS) [3–8]. In a prospective cohort study, our group observed that 46% of the infants that were born to ZIKV-infected mothers bore abnormal clinical or brain imaging findings, including four infants with microcephaly, regardless of the trimester in pregnancy [9]. Indeed, in a little while, a case-control study confirmed the association between the infection and CZS, and ZIKV epidemic was declared a public health emergence of international concern [3,10]. Nevertheless, not all infants that are born to ZIKV infected mothers will develop CZS, and it is not clear what maternal and/or fetal factors contribute to infant adverse neurologic outcomes. One important risk factor for CSZ is infection within the first trimester of pregnancy, which poses almost twice as high a risk of severe outcomes such as CNS abnormalities when compared with third trimester infections [9]. Furthermore, maternal nutritional and social factors, such as consumption of improper water and poor protein diet, have been related to CZS development [11,12]. Although these environmental factors do not completely explain CZS outcomes, it has been reported that genetic background can influence these outcomes. Thus, studies with a smaller cohort utilizing high throughput sequencing techniques independently identified maternal adenylate cyclase and newborn collagen-encoding genes associated with abnormal outcomes due to ZIKV infection during pregnancy [13–15].

During other congenital infections, namely TORCH (Toxoplasma, Others, Rubella, Cytomegalovirus, Herpes), which may cause malformations, placenta has been described as playing a crucial role in mother to fetus transmission [16]. In zika, one of the hypotheses for the emergence of adverse neurological outcomes is that ZIKV can infect and cross trophoblast cell layers as cargo, ultimately reaching the fetal neurologic system and causing direct damage. On the other hand, ZIKV infection causes an innate immunological imbalance, excessive inflammation and vascular permeability dysfunction in the placenta, which may contribute to disrupting embryonic brain development [17–24].

Interferons (IFN) are key players of the innate immune response against viral infection, inducing hundreds of interferon-stimulated genes (ISGs) that act directly against virus components [25]. Among these ISGs, ubiquitin-like protein ISG15, induced by type I IFNs, is one of the most strongly and rapidly induced, inhibiting viral replication and modulating host immunity [26–28]. Another ISG, IFN-induced protein with tetratricopeptide repeats 5, IFIT5 (ISG58), activates IRF3/NF-κB pathway, which induces higher type I IFNs and proinflammatory mediators [29]. It has been described that ZIKV disrupts type I IFN, harming phosphorylation of STAT1 and STAT2 [30,31]. In addition to a major role in antiviral defense, an exacerbated type I IFN response was demonstrated to be threatening for newborn development [32], indicating that a balanced production of type I IFNs could be effective in controlling infection and inflammation. Type III interferons (a.k.a. IFN-λ 1-4) present augmented expression during ZIKV infection in susceptible placental cells and higher levels of IFN-λ antagonize type I IFNs [21,33–35]. In this regard, administration of exogenous IFN-λ in mice led to signatures with balanced expression of ISGs (*IFI44L*, *OASL*, *OAS1*, and *MX1*) and inhibition of ZIKV replication, suggesting a therapeutic potential [36,37].SNPs in the vicinity of *IFNAR1* and *IFNL1-4* loci have been associated with outcomes of viral infections, such as hepatitis B and C [38–40]. Variants within *IFNAR1* have been associated with an error of innate immunity related to severe viscerotropic adverse events following vaccination with another flavivirus: attenuated yellow fever virus [41]. The *IFNL4* rs12979860 CC genotype has been associated with persistent low levels of ISGs *IFIT1*, *IFIT2*, *IFIT3*, and *OAS1* in postpartum normal pregnancy [42]. In another flavivirus infection, hepatitis C, rs12979860 CC is a marker for good prognosis in chronic patients treated with IFNα and ribavirin [43,44].

In this paper, we describe the association between the genetic background of newborns and mothers from ZIKV infected pregnancy and CZS development, focusing on SNPs in *IFNAR1*, *IFNL2* and *IFNL4* loci, as well as the functional consequences of specific genotypes for the immunological imbalance in at term placentas from Zika pregnancy cases.

## 2. Materials and methods

### Human subjects and sample collection

Our studies made use of the ongoing prospective clinical cohort study of ZIKV+ pregnant women and their infants at a maternal and child hospital (IFF/Fiocruz) in Rio de Janeiro, Brazil (IRB/CAAE: 52675616.0.000.5269). In this cohort, pregnant women who were ZIKV+ received their prenatal care at IFF/Fiocruz. Since December 2015, a total of 301 mothers who were suspected of having been infected by ZIKV during gestation were referred to IFF — a major public reference hospital in Rio de Janeiro for congenital infections and malformations. Here, we utilized a subpopulation of 143 newborns and 153 mothers from the IFF cohort, including 3 pairs of bi-chorionic and bi-amniotic twins. Among the newborns, 66 (46%) were considered symptomatic, presenting CNS (Central Nervous System) and/or eye abnormalities at birth. CNS abnormalities include microcephaly, ventriculomegaly, cerebral calcifications, posterior fossa abnormalities, pachygyria, and lissencephaly, which were not mutually exclusive. Eye abnormalities included optic nerve atrophy, chorioretinal atrophy, pigment mottling and hemorrhage, which occurs frequently and is associated with CNS alterations. From those cases, 84 placentas (74 from congenital ZIKV infections and 10 from uninfected patients) were accessed, processed and analyzed for gene expression. Samples from mothers were tested for HIV, evidence of past Dengue virus (DENV) infection (by DENV IgG and IgM), and Chikungunya virus (CHIKV) (blood PCR). Maternal demographic, medical/prenatal history and clinical findings were entered into case-report forms. All infants underwent routine clinical and extensive neurologic evaluation at the time of birth and were tested for CHIKV infection (blood PCR), syphilis and TORCH infections (toxoplasmosis, rubella, CMV, and herpes simplex virus; as determined by standard testing). Infants were evaluated for the following adverse neurologic outcomes: (a) microcephaly (head-circumference z score of less than −2), (b) abnormal brain imaging by pre- or post-natal ultrasound (e.g., computed tomography and/or magnetic resonance imaging), and/or (c) abnormal clinical examination (including neurologic, ocular, and/or auditory with abnormalities confirmed by a multidisciplinary team of neonatologists, neurologists, infectious disease specialists, geneticists, ophthalmologists, and physical therapists). From this cohort, adverse clinical outcomes (mostly neurologic) were reported at birth in 41% of infants who were born to mothers during all trimesters. Our study included ZIKV+ pregnant adult women >18 years of age and their infants. Exclusion criteria included maternal HIV infection and pregnancies complicated by other congenital infections, known to cause infant neurologic damage (e.g., TORCH, CHIKV). Placental samples were collected at the time of delivery from the umbilical cord insertion region and stored in RNA later until RNA extraction. For DNA analysis, 5 mL of m blood was collected from pregnant women at study enrollment and an oral swab was collected from newborns.

### Genetic studies: SNP selection and linkage disequilibrium analysis

Selection of candidate SNPs for the case-control association study was performed by integrating different tools: Principal Component Analysis (PCA), ANNOVAR [45], allele frequencies, literature and HAPLOVIEW [46]. First, all SNPs located in the *IFNL* (chr19:39,733,272-39,736,609-GRCh37/hg19) and *IFNAR1* regions (chr21:34,696,734-34,732,168-GRCh37/hg19) were recovered from African (ENS, GWD, LWK, MSL, and YRI) and European (CEU, FIN, GBR, IBS, and TSI) populations from phase 3 of the 1000 Genomes Project [47]. Then, Principal Component Analysis (PCA) was performed using EIGENSOFT4.2 [46]. The use of this strategy in the selection of functional SNPs assumes that, since the analyzed variability is of a functional genome region (meaning: a gene), the clusters generated by PCA would be mainly influenced by functionality. Thus, SNPs with high weight for principal component 1 (PC1) could be potential candidates for having a functional role. SNPs were thus sorted by decreasing values of “SNP weight” for PC1, and functional annotation of all SNPs was performed using ANNOVAR [45], with ref Gene hg19 (11 Dez 2015). According to the functional category identified by ANNOVAR, “SNP weight” for PC1 (with SNP weight values within the highest 30, called top SNPs), minimal allele frequencies (MAF) in African and European populations (> 0.1) and associations with infectious diseases already reported in the literature, SNPs present in the *IFNL* region were selected for genotyping and haplotype construction. Haplotype inferences using selected SNPs, haplotype frequencies and linkage disequilibrium (LD) analysis for all studied populations were performed using HAPLOVIEW [46]. To select SNPs, PCA was used to retrieve those located either in the *IFNAR1* or *IFNL* regions, which are found among the African and European populations from the 1000 Genomes Project (Supplementary Fig. 1A). Within *IFNL*, we selected four representative SNPs (rs12979860, rs4803222, rs8109886, and rs8099917). In the *IFNAR1* region, SNPs rs2843710, rs2257167, rs17875834, rs2834202 were selected to construct the haplotypes in parental populations. Allele frequencies, annotation, and reference of the selected SNPs are described in Supplementary Table 1. Linkage disequilibrium (LD) analysis and haplotype arrangements indicated ancestry-specific patterns for these two genomic regions. (Supplementary Fig. 1B). The *IFNAR1* arranged haplotypes suggest rs2843710, rs2257167 are tags to discriminate Europeans and Africans, while *IFNL* rs12979860 and rs8109886 SNPs also present very different frequencies among the major Brazilian parental populations (Supplementary Table 2).

**Table 1.** Association study with newborn *IFNAR1* and *IFNL* SNPs and CZS abnormalities. DNA samples from 143 newborns with congenital infection by ZIKV, with or without CZS, were genotyped for *IFNAR1* SNPs rs2257167 and rs2843710 and *IFNL* SNPs rs8109886, rs12979860 rs8099917, and rs4803222. The total number of genotyped samples for each SNP may vary due to genotype miscalling. Major allele was used as baseline. Odds ratio (OR) with 95% confidence interval (CI) and p-values. We conducted type I error adjustment of multiple comparisons by the False Discovery Rate (FDR) method.

|  | No CZS findings |  | Abnormal CZS |  | Adjusted by trimester of ZIKV exposition and ancestry |  |  |  |  |
| --- | --- | --- | --- | --- | --- | --- | --- | --- | --- |
|  | N<sup>a</sup> | % | N<sup>a</sup> | % | OR | lower | upper | p-value | FDR* p-value |
| rs2257167 |  |  |  |  |  |  |  |  |  |
| G/G | 48 | 77.4 | 32 | 57.1 | Ref |  |  | 0.026 | 0.04324 |
| C/G | 12 | 19.4 | 21 | 37.5 | 3.51 | 1.22 | 10.13 |  |  |
| C/C | 2 | 3.2 | 3 | 5.4 | 5.06 | 0.66 | 38.54 |  |  |
| G/G | 48 | 77.4 | 32 | 57.1 | Ref |  |  | 0.0073 | 0.02646 |
| C/G-C/C | 14 | 22.6 | 24 | 42.9 | 3.73 | 1.36 | 10.21 |  |  |
| G/G-C/G | 60 | 96.8 | 53 | 94.6 | Ref |  |  | 0.2321 | 0.27860 |
| C/C | 2 | 3.2 | 3 | 5.4 | 3.27 | 0.46 | 23.45 |  |  |
| G/G-C/C | 50 | 80.6 | 35 | 62.5 | Ref |  |  | 0.0288 | 0.04324 |
| C/G | 12 | 19.4 | 21 | 37.5 | 3.02 | 1.08 | 8.40 |  |  |
| Additive | 62 | 52.5 | 56 | 47.5 | 2.80 | 1.25 | 6.29 | 0.0088 | 0.02646 |
| rs2843710 |  |  |  |  |  |  |  |  |  |
| G/G | 23 | 39.7 | 22 | 37.9 | Ref |  |  | 0.2658 | 0.53150 |
| C/G | 28 | 48.3 | 23 | 39.7 | 1.00 | 0.42 | 2.38 |  |  |
| C/C | 7 | 12.1 | 13 | 22.4 | 2.38 | 0.74 | 7.72 |  |  |
| C/C | 23 | 39.7 | 22 | 37.9 | Ref |  |  | 0.5694 | 0.68322 |
| C/G-G/G | 35 | 60.3 | 36 | 62.1 | 1.26 | 0.56 | 2.85 |  |  |
| C/C-C/G | 51 | 87.9 | 45 | 77.6 | Ref |  |  | 0.1035 | 0.53150 |
| G/G | 7 | 12.1 | 13 | 22.4 | 2.39 | 0.82 | 6.99 |  |  |
| C/C-G/G | 30 | 51.7 | 35 | 60.3 | Ref |  |  | 0.4930 | 0.68322 |
| C/G | 28 | 48.3 | 23 | 39.7 | 0.76 | 0.35 | 1.67 |  |  |
| Additive | 58 | 50 | 58 | 50 | 1.42 | 0.81 | 2.48 | 0.2115 | 0.53150 |
| rs12979860 |  |  |  |  |  |  |  |  |  |
| C/C | 31 | 40.8 | 22 | 33.3 | Ref |  |  | 0.5803 | 0.87045 |
| C/T | 33 | 43.4 | 28 | 42.4 | 1.36 | 0.56 | 3.32 |  |  |
| T/T | 12 | 15.8 | 16 | 24.2 | 1.81 | 0.57 | 5.76 |  |  |
| C/C | 31 | 40.8 | 22 | 33.3 | Ref |  |  | 0.3585 | 0.85948 |
| C/T-T/T | 45 | 59.2 | 44 | 66.7 | 1.48 | 0.64 | 3.4 |  |  |
| C/C-C/T | 64 | 84.2 | 50 | 75.8 | Ref |  |  | 0.4297 | 0.85948 |
| T/T | 12 | 15.8 | 16 | 24.2 | 1.52 | 0.53 | 4.33 |  |  |
| C/C-T/T | 43 | 56.6 | 38 | 57.6 | Ref |  |  | 0.7835 | 0.94016 |
| C/T | 33 | 43.4 | 28 | 42.4 | 1.12 | 0.5 | 2.5 |  |  |
| Additive | 76 | 53.5 | 66 | 46.5 | 1.35 | 0.77 | 2.37 | 0.2971 | 0.85948 |
| rs8099917 |  |  |  |  |  |  |  |  |  |
| T/T | 57 | 75 | 43 | 66.2 | Ref |  |  | 0.3497 | 0.52456 |
| G/T | 17 | 22.4 | 18 | 27.7 | 2.05 | 0.76 | 5.5 |  |  |
| G/G | 2 | 2.6 | 4 | 6.2 | 1.31 | 0.2 | 8.61 |  |  |
| T/T | 57 | 75 | 43 | 66.2 | Ref |  |  | 0.1661 | 0.49821 |
| G/T-G/G | 19 | 25 | 22 | 33.8 | 1.89 | 0.76 | 4.71 |  |  |
| T/T-G/T | 74 | 97.4 | 61 | 93.8 | Ref |  |  | 0.9071 | 1 |
| G/G | 2 | 2.6 | 4 | 6.2 | 1.12 | 0.17 | 7.25 |  |  |
| T/T-G/G | 59 | 77.6 | 47 | 72.3 | Ref |  |  | 0.1551 | 0.49821 |
| G/T | 17 | 22.4 | 18 | 27.7 | 2.02 | 0.76 | 5.4 |  |  |
| Additive | 76 | 53.9 | 65 | 46.1 | 1.53 | 0.73 | 3.22 | 0.2491 | 0.49822 |
| rs8109886 |  |  |  |  |  |  |  |  |  |
| A/A | 20 | 28.6 | 21 | 32.8 | Ref |  |  | 0.8777 | 1 |
| C/A | 34 | 48.6 | 30 | 46.9 | 0.92 | 0.37 | 2.31 |  |  |
| C/C | 16 | 22.9 | 13 | 20.3 | 0.74 | 0.23 | 2.36 |  |  |
| A/A | 20 | 28.6 | 21 | 32.8 | Ref |  |  | 0.7462 | 1 |
| C/A-C/C | 50 | 71.4 | 43 | 67.2 | 0.87 | 0.36 | 2.07 |  |  |
| A/A-C/A | 54 | 77.1 | 51 | 79.7 | Ref |  |  | 0.6323 | 1 |
| C/C | 16 | 22.9 | 13 | 20.3 | 0.78 | 0.29 | 2.12 |  |  |
| A/A-C/C | 36 | 51.4 | 34 | 53.1 | Ref |  |  | 0.9291 | 1 |
| C/A | 34 | 48.6 | 30 | 46.9 | 1.04 | 0.47 | 2.3 |  |  |
| Additive | 70 | 52.2 | 64 | 47.8 | 0.87 | 0.49 | 1.54 | 0.6267 | 1 |
| rs4803222 |  |  |  |  |  |  |  |  |  |
| A/A | 30 | 42.9 | 31 | 51.7 | Ref |  |  | 0.5137 | 0.71400 |
| C/A | 35 | 50 | 23 | 38.3 | 0.64 | 0.29 | 1.39 |  |  |
| C/C | 5 | 7.1 | 6 | 10 | 0.93 | 0.23 | 3.77 |  |  |
| A/A | 30 | 42.9 | 31 | 51.7 | Ref |  |  | 0.3049 | 0.71400 |
| C/A-C/C | 40 | 57.1 | 29 | 48.3 | 0.68 | 0.32 | 1.43 |  |  |
| A/A-C/A | 65 | 92.9 | 54 | 90 | Ref |  |  | 0.8327 | 0.71400 |
| C/C | 5 | 7.1 | 6 | 10 | 1.16 | 0.3 | 4.46 |  |  |
| A/A-C/C | 35 | 50 | 37 | 61.7 | Ref |  |  | 0.2504 | 0.71400 |
| C/A | 35 | 50 | 23 | 38.3 | 0.64 | 0.3 | 1.37 |  |  |
| Additive | 70 | 53.8 | 60 | 46.2 | 0.81 | 0.45 | 1.46 | 0.4768 | 0.71400 |

### Genomic DNA extraction and SNP genotyping analysis

DNA extraction was performed from saliva swabs or whole blood cells collected from each individual newborns (n=143) and mothers (n=153), respectively, using the salting out method. Following extraction, DNA was quantified with a Nanodrop ND 1000 spectrophotometer (Nanodrop Technologies). After PCA, LD and haplotype analysis, the following tag polymorphisms were genotyped due to their representativeness within the corresponding genomic regions: *IFNL*2-*IFNL*4: rs8099917 (C11710096_10) located 8.9 kb upstream of the *IFNL*4 (T > G) start codon and rs8109886 (C11710100_10) located 3.3 kb upstream of the *IFNL*4 (A > C) start codon; *IFNL*4: rs12979860 (C7820464_10) in intron 4 (C > T) and rs4803222 (C7820457_10) in the 5’ UTR (C > G); *IFNAR1*: rs2257167 (C 16076297_10) located within exon 4 of *IFNAR1* (G>C) and rs2843710 (C 26796048_10) located within the promoter region of *IFNAR1* (C > G). All SNPs were genotyped using the allelic discrimination method for real-time TaqMan assays (Applied Biosystems) using either the ABI Prism 7000 Sequence Detection System or the Step One Plus Real-time PCR System. Approximately 50 ng of DNA was used in the genotyping reaction. Statistical analyses were performed using “snpassoc”, “genetics” and “haplo.stats” packages in software R version 2.11.1, as previously described [48]. Briefly, genotype frequencies were tested for Hardy–Weinberg equilibrium (HWE) using a Chi-square test. The genotypic, allelic, and carrier frequencies were calculated and compared in cases and controls by conditional logistic regression adjusted for ancestry and trimester of infection. Next, we compared the frequencies between CZS and no CZS, separately. Linkage disequilibrium values for SNPs studied in *IFNL* were estimated by r^2^ and haplotype frequencies were compared between cases and controls by logistic regression, also adjusted for ancestry and trimester of infection. For mother-child SNP interaction, we used EMIM analysis using a multinomial model to test the existence (and estimate) of genotype relative risk parameters that may increase (or decrease) the possibility that a child is affected, as described previously [49].

#### Ancestry analysis

Since the Brazilian population is highly admixed and ethnic classification is not uniformly defined, ancestry data is necessary to adjust the logistic regression and eliminate bias in genetic associations [50]. Thus, DNA samples were genotyped for 46 Ancestry Informative Markers (AIM)-Indels in a multiplex PCR system followed by capillary electrophoresis in an ABI 3500 Genetic Analyzer (Thermo Fisher), as described previously [51,52]. Allele calls were obtained by GeneMapper v.4.1 and results for individual and global ancestry estimates were performed by using the HGDP-CEPH diversity panel as a reference (European, African and Native-American; K=3) in STRUCTURE v2.3. In the logistic regression performed in R, covariates AFR+EUR were used to control for population stratification along with trimester of infection.

### ZIKV PCR detection

RT-qPCR was performed using the 2x QuantiTect Probe RT-PCR kit (Qiagen, Valencia, CA, USA) with the same primers and cycle times as previously described[53]. All the assays were carried out in triplicate and fluorescence curves that crossed the threshold within or below 38 cycles were considered positive.

### Gene expression profile analysis

Analysis of gene expression in placental tissue from pregnant mothers (control, with or without CZS samples) was performed using Fluidigm (Biomark platform) assays. Detailed data available under request. Our experimental design followed a previously described workflow [54].

### Real-time RT-PCR expression analysis

From routines created in R for parsing raw foreground and background intensities, exported from the commercial platform Fluidigm®, we carried out background correction and exploratory data analysis: fluorescence accumulation and melting curve graphs of Rn for each reaction with each gene. For relative quantification of expression, the fluorescence accumulation data of each sample were used for fitting four parameter sigmoid curves using the qPCR^56^ library from R statistical package version 3.4.1[48]. For each amplification, the cycle of quantification was determined as the maximum of the second derivative of the fit sigmoid curve and the efficiency, as the ratio between the fluorescence of the cycle of quantification and the fluorescence of the cycle that immediately preceded that. For each gene, efficiency was estimated by the mean of all the efficiencies for each amplification reaction for that gene. Endogenous controls used for normalizing between different amplified samples were selected by the geNorm method. Normalization factors were estimated for each sample using the geometric average of the selected normalized genes [56].

### Statistical analysis of gene expression

Pairwise comparisons of log-transformed (base 2) normalized expression means between/among groups of interest were performed by contrasts/differences (fold-changes) obtained after both bi- and multivariate linear models were modified by ordinary least square regressions. Whenever the variable of interest had more than two levels, p-values were corrected by the Tukey Honest Significant Difference post-Hoc method [57]. After gene-per-gene pairwise comparisons, we conducted a Type I error adjustment for multiple comparisons by the Holm-Bonferroni method [58]. Different sets of confounding variables were selected by clinical experts and included in the multivariate models to adjust the fit effects for different variables of interest for all genes. For the analysis, two-tailed levels of significance ≤ 0.01, 0.05, and 0.1 were considered as “highly significant,” “significant,” and “suggestive,” respectively.

## 3. Results

### 3.1. Newborn *IFNAR1* rs2257167 are associated with CZS outcome

The DNA samples from whole blood of 143 newborns and 153 mothers from ZIKV-infected pregnancies, with development of abnormal CZS (cases) or otherwise (controls), were genotyped for SNPs encompassing *IFNAR1* (rs2257167 and rs2843710) and *IFNL4* genes (rs12979860 and rs4803222), and within *IFNL2* and *IFNL4* genes (rs8099917 and rs8109886). The frequency of each SNP was verified in cases and controls and CZS outcome was evaluated. Genotype frequencies were found to be in HWE for all SNPs tested. Data were adjusted by genetic ancestry and the trimester of pregnancy in which ZIKV infection occurred (when symptoms of ZIKV infection were detected). CZS risk was observed for CG/CC carriers of SNP rs2257167 following FDR correction, OR = 3.73; CI = 1.36-10.21, *Pcorrected* = 0.0264 (Table 1). No significant differences were observed in the frequencies between cases and controls in any other SNP tested. Other analyses including *IFNAR1* and *IFNL* haplotypes, and mother genotypes did not show any significant results (Supplementary Tables 3, 4, and 5).

### 3.2. *IFNAR1* rs2257167 CG/CC genotypes are CZS risk factors in ZIKV infection during second and third trimester of pregnancy

Following previous clinical studies [9], in our cohort, the determination of the trimester of pregnancy in which ZIKV infection occurs was a strong predicting factor for CZS outcome, along with *IFNAR1* genotype (Fig. 1). ZIKV infections during the first trimester of pregnancy culminate with 64.1% of cases in CZS, more than twice the chance of CZS development in the second (19.2%) or third (30.3%) trimesters. Interestingly, data showed that newborns with rs2257167 CG/CC genotypes presented higher frequencies of CZS considering all pregnancy periods, although they presented twice the frequency compared with newborns with the GG genotype in second (16.6%-GG *vs* 41.6% −CG/CC) and third (18.75%-GG *vs* 50%-CG/CC) trimesters.

**Fig 1.**
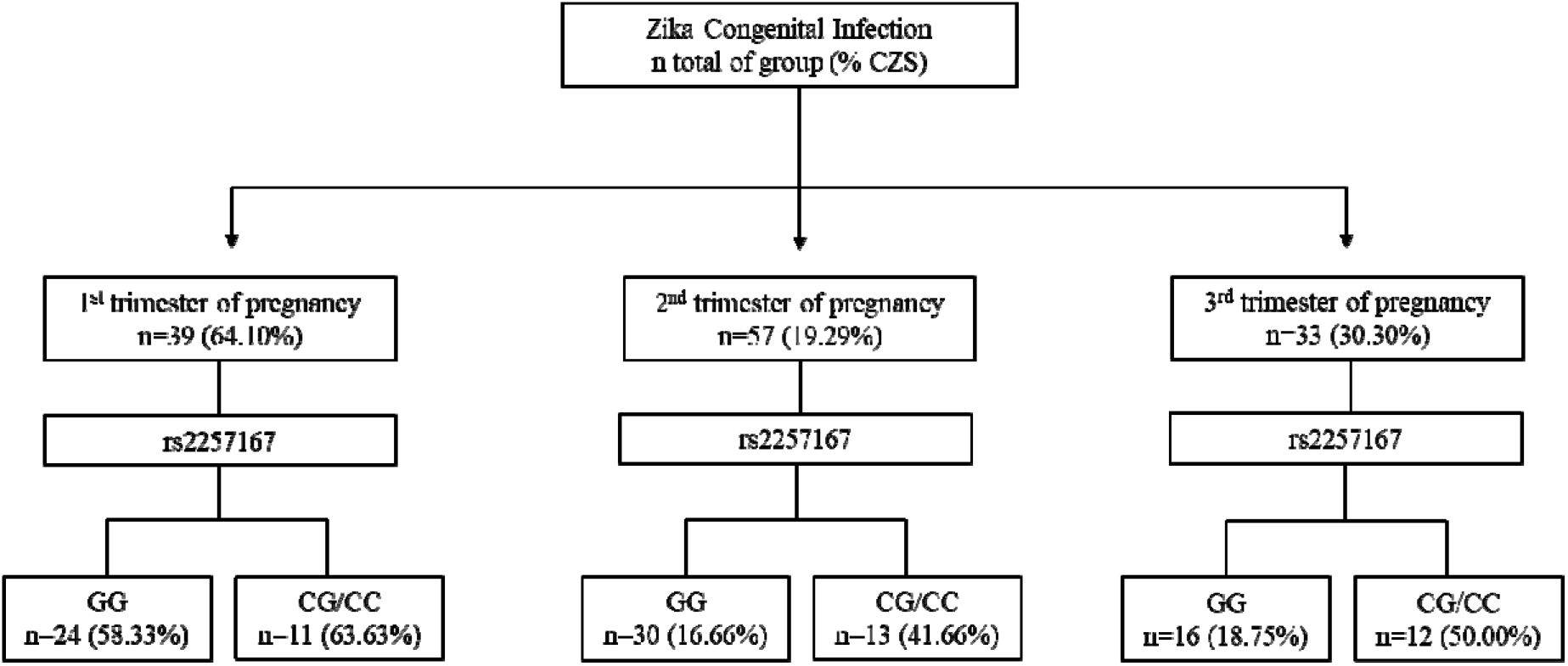
Event-based flowchart of CZS occurrence. The trimester of pregnancy in which first Zika symptoms occur and newborn genotypes of rs2257167 were used as independent variables to determine the association with CZS outcome. Absolute number of newborns per group (n) and CSZ percentage are shown. Concerning infections in the first trimester of pregnancy, we found that irrespective of rs2257167 genotypes, 64.10% or more of the cases developed CZS, in contrast with second and third trimesters. Considering rs2257167, newborn CG/CC genotype seems to be associated with CZS risk in second and third trimesters.

### 3.3. Congenital ZIKV infection leads to an immunological imbalance in placenta

To functionally verify how ZIKV could influence severe congenital outcomes across associated *IFNAR1* genotypes, we performed gene expression analyses from the placental tissues, most of which were fetal, obtained at time of delivery from 10 uninfected pregnant woman and 74 congenital ZIKV cases (Supplementary Table 6). We used a multiplex RT-qPCR analysis of 96 candidate immunity-associated genes and only statistically significant results are presented. First, all ZIKV RT-PCR positive samples at term showed higher gene expression of most genes analyzed. ZIKV congenital infections occurring in the third trimester of pregnancy resulted in highest gene expression levels. Mothers exhibiting ≥ 40 years of age (y/o) expressed lower levels of inflammatory genes, suggesting natural immunological senescence (Supplementary Table 7). Because of these intrinsic differential expression profiles, ZIKV RT-qPCR positive placenta, trimester of exposure to ZIKV, and mothers’ age (≥ 40 y/o) were considered as variables in gene expression analysis of congenital ZIKV cases.

Comparing placental gene expression from congenital ZIKV infections *vs* uninfected pregnant women results showed that ZIKV leads to a typical inflammatory response in placenta including higher expression of: anti-viral type I IFN genes (*IFIT5*, *IFNA1*, and *IFNB*), type II interferon (*IFI16*), cytokine signaling (*IL22RA* and *IP10*), and interferon regulatory factors (*IRF7* and *IRF9*); together with decreased expression of *TYRO3* (Fig. 2).

**Fig 2.**
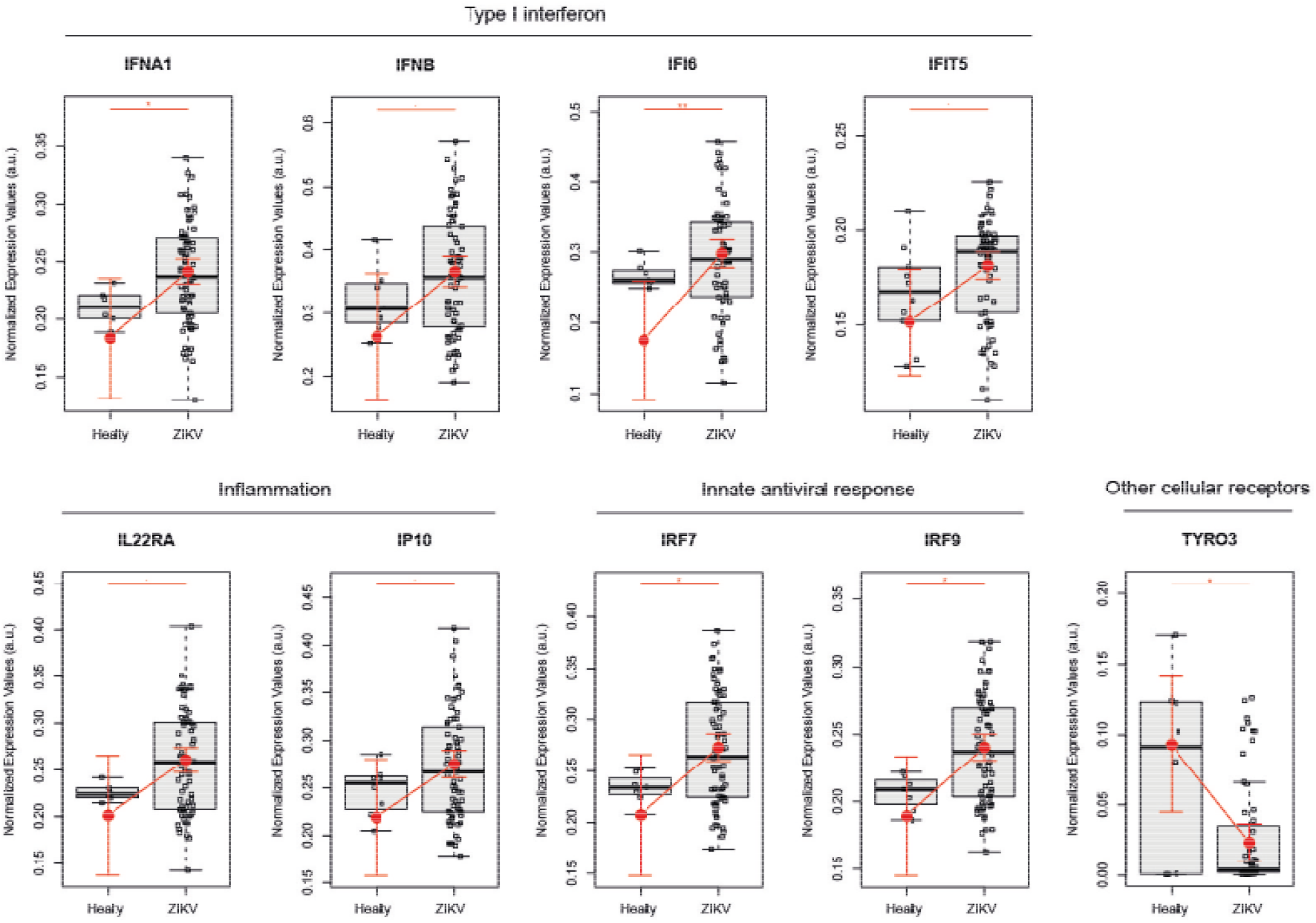
ZIKV infection leads to an immunological imbalance in placenta. Placental gene expression profile in healthy (N = 10) versus ZIKV-infected women (N = 74). Each dot corresponds to one placenta analyzed. The number of dots varies according to gene analyzed due to failed amplifications. Median and standard deviation of gene expression values are normalized by the housekeeping genes selected by the geNorm and NormFinder as well as *18S* ribosomal RNA and *RLP13* ribosomal protein L13 (grey boxes). Values are adjusted by mothers’ age (below or equal to/above 40 years of age) and trimester of infection (the trimester of pregnancy in which the first Zika symptoms occur or asymptomatic ZIKV infections) (red lines). Pairwise comparisons of log-transformed (base 2) normalized expression means between/among the groups of interest were performed by contrasts/differences (fold-changes), obtained after both bi- and multivariate linear models were modified by ordinary least square regressions. Whenever the variable of interest had more than two levels, p-values were corrected by the Tukey Honest Significant Difference post-Hoc method. After gene-per-gene pairwise comparisons were made, we conducted a Type I error adjustment for the multiple comparisons by the Holm-Bonferroni method. P-values ** ≤ 0.01, * ≤ 0.05 and ≤ 0.1.

Interestingly, clustering results by ZIKV PCR detection in at term placentas, ZIKV PCR+ placentas showed increased expression of type I and III interferon pathway genes analyzed, with exception to *IFITM1*, which presented reduced expression (Supplementary Figure 2). Data also demonstrated augmented expression of *CLEC5A* and *DCSIGN* as well as *TLR3* and *TLR8* pattern-recognition receptors, *BCL2*, *CARD9* (apoptosis-related), *IL18* and *AIM2* (type II interferon) and *IFNL1* (type III interferon). Corroborating the ZIKV-infection pro-inflammatory placental profile, results showed elevated expression of chemokine-related genes (i.e., *CCR2*, *CCR3*, *CCR5*, *MIP1A*, and *IP10*) and other cytokine-related genes (i.e., *IL22A*, *MMP2*, and *TNF*) besides *NRPL3*, which denotes the presence of inflammasome activation in ZIKV PCR+ placental samples. Results also showed an increase in the expression of *IL10* (Supplementary Fig. 2).

### 3.4. Decreased *IFNL2* and augmented type I IFN in placenta at term is associated with newborn CZS abnormalities

Next, we tested whether gene expression signatures of placentas from ZIKV-infected women could be associated with the presence or absence of abnormal CZS. These analyses illustrated *IFNL2* and *ISG15* significant decreases in newborns with CZS. On the other hand, *IFIT5* increased significantly in newborns in the CZS group (Fig. 3).

**Fig 3.**
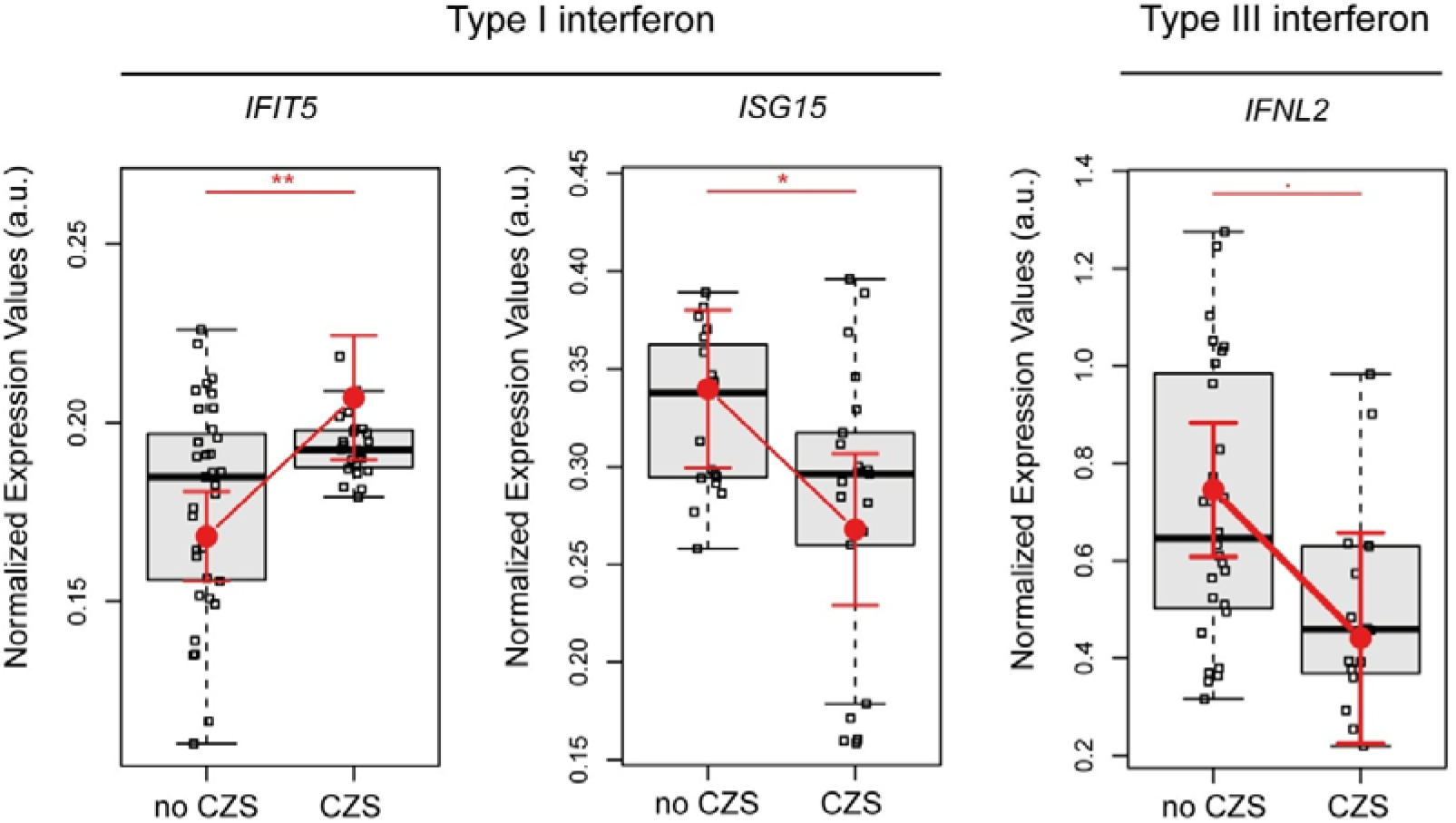
Placental gene expression associated with CZS. Detailed graphs of differentially expressed genes in placenta without CZS (No CZS; N = 45) or with abnormal CZS (CZS; N = 29). Each dot corresponds to one placenta analyzed. The number of dots varies according to gene analyzed due to failed amplifications. Median and standard deviation of gene expression values are normalized by housekeeping genes selected by geNorm and NormFinder as well as *18S* ribosomal RNA and *RLP13* ribosomal protein L13 (grey boxes). Values are adjusted by mothers’ age (below or equal to/above 40 years of age) and infection trimester (trimester of pregnancy in which the first Zika symptoms occur or asymptomatic ZIKV infections) (red lines). Pairwise comparisons of log-transformed (base 2) normalized expression means between/among the groups of interest were performed by contrasts/differences (fold-changes), obtained after both bi- and multivariate linear models were modified by ordinary least square regressions. Whenever the variable of interest had more than two levels, p-values were corrected by the Tukey Honest Significant Difference post-Hoc method. After gene-per-gene pairwise comparisons were carried out, we conducted a Type I error adjustment for multiple comparisons by the Holm-Bonferroni method. P-values ** ≤ 0.01, * ≤ 0.05, and ≤0.1.

### 3.5. Genotypes rs2257167 CG/CC are associated with increased placental type I IFN and inflammatory response

We clustered 33 newborns and 34 mothers according to GG or CG/CC genotypes of rs2257167 to assess how *IFNAR1* newborn background influences the placental gene expression profile. Placentas from rs2257167 CG/GG newborns showed significantly increased expression of *IFIT5* and genes related with the inflammatory response (*IL8, IL23A, MMP9, MIP1A, MARCO, NRLP1, and TNFSF15*) (Fig. 4A). Given that ZIKV PCR positive placenta present a more active immunological response, we clustered placenta expression by ZIKV PCR and rs2257167 genotypes. Considering ZIKV PCR positive placenta, we observed that CG/CC newborns presented augmented expression of *IFNAR1* and genes related with inflammation (*IL8* and *TNF*), innate antiviral response (*DAP12*, *RIPK2*, *SOD2*, and *STAT2*), toll like receptors (*TLR2*, and *TLR4*), *CD36* and *TYRO3* (Fig 4B).

**Fig 4.**
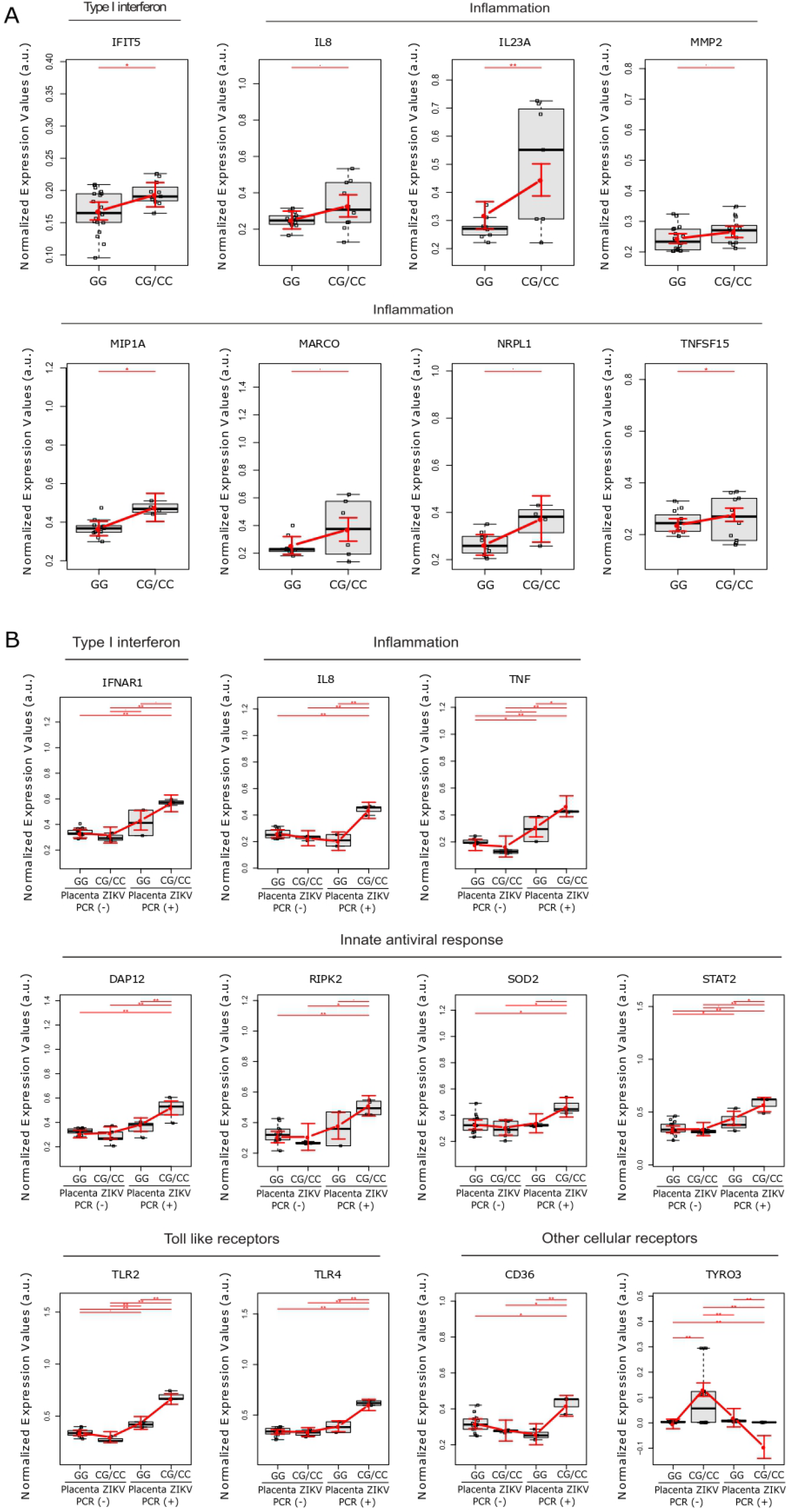
Placental gene expression is modulated by newborn rs2257167 genotypes in ZIKV infected pregnancy. (A) Detailed graphs of differentially expressed genes in placenta from rs2257167 GG (N = 20) and CG/CC (N = 13) newborns. Each dot corresponds to one placenta analyzed. The number of dots varies according to gene analyzed due to failed amplifications. Median and standard deviation of gene expression values are normalized by housekeeping genes selected by geNorm and NormFinder as well as *18S* ribosomal RNA and *RLP13* ribosomal protein L13 (grey boxes). Values are adjusted by mothers’ age (below or equal to/above 40 years of age) and infection trimester (trimester of pregnancy in which first Zika symptoms or asymptomatic ZIKV infections occur) (red lines). Pairwise comparisons of log-transformed (base 2) normalized expression means between/among groups of interest were performed by contrasts/differences (fold-changes), obtained after both bi- and multivariate linear models were modified by ordinary least square regressions. Whenever the variable of interest had more than two levels, p-values were corrected by the Tukey Honest Significant Difference post-Hoc method. After gene-per-gene pairwise comparisons were made, we conducted a Type I error adjustment for the multiple comparisons by the Holm-Bonferroni method. P-values**≤ 0.01, * ≤ 0.05, and ≤ 0.1.

## 4. Discussion

Despite the high risk, ZIKV infection during pregnancy is necessary, albeit not enough, to induce CZS. We hypothesized whether host genetic background, especially SNPs in *IFNAR1* and *IFNL*, contributes to CSZ development and conducted a case-control study with CZS cases and healthy ZIKV^+^ mothers. Specifically, our results demonstrated that newborns who carried the CG/CC genotypes of SNP rs2257167 (*IFNAR1*) had a 3.7 higher risk of developing abnormal CZS, resulting from ZIKV infection during pregnancy, compared to those with the GG genotype. The rs2257167 polymorphism is a missense val(G)>leu(C) SNP that can potentially alter the structure of the protein [59]. In chronic hepatitis B infected patients, the presence of the C allele was associated with higher plasma levels of the aspartate and alanine amino-transferase hepatic enzymes [60]. Another study among chronic HBV-infected patients suggested that patients carrying the rs2257167 CC (leu/leu) genotype presented higher expression levels of *IFNAR1* in PBMCs when compared with patients carrying the GG genotype (val/val)[59].

Our data also highlights the immunomodulatory role of rs2257167 and how this SNP influences CZS frequency, especially when ZIKV infections occur in second and third trimesters of pregnancy. This data also shows the importance of *IFNAR1* genetic background regulating placental gene expression culminating in CZS development. Studies using mice models and *ex vivo* placental cultures demonstrated that regions and maturity of placentas will provide different responses against ZIKV [37,61]. Generally, fetal-derived tissues developed from midgestational placenta are more restrictive to ZIKV replication [61]. In fact, *in vitro* cultures show that ZIKV possesses high tropism for trophoblasts from the first trimester of pregnancy [62]. Altogether, these data corroborate our findings, considering that ZIKV faces an IFN immunological barrier in midgestational or older placentas, while rs2257167 CG/CC carriers with higher type I and lower type III IFNs would unbalance type I / type III IFNs towards a pronounced and exacerbated type I IFN production leading to CZS susceptibility.

Indeed, a high type I IFN expression phenotypic pattern in rs2257167 CG/CC individuals was observed suggesting they cannot efficiently regulate exacerbated type I IFN, which might be one of the factors leading to CZS. Notably, placenta from the abnormal CZS cases is associated with an increased expression of *IFIT5*, which is an important enhancer of type I IFN and a proinflammatory response [29]. This profile of augmented type I IFN associated with severity is corroborated in a ZIKV infected mice model [63] and other TORCH infections [64,65]. However, it is noteworthy that studies in mice models demonstrated that the lack of a type I IFN response also lead to CSZ [63,66] indicating that only optimal levels of type I IFN could possibly confer a healthy pregnancy upon Zika and probably other congenital infections. Notwithstanding, positive ZIKV PCR in at term placenta highlights aspects of ZIKV placental infection as well as high *TLR3* and *TLR8* mRNA expression, which could contribute as pattern recognition receptors for ZIKV uptake [67–70]. Although *in vitro* studies strongly suggest that *TYRO3* is the main entry receptor for ZIKV [67,71], persistent ZIKV-infected placentas showed a decreased expression of *TYRO3* in ZIKV PCR+ individuals, corroborating recent findings in mice indicating that in complex organisms these receptors do not appear to be required for ZIKV infection [72]. Otherwise, increased expression of *CLEC5A* and *DCSIGN* could also suggest alternative routes for virus uptake or antiviral cellular response, as reported for other flavivirus infections [73–76]. Higher expression of *BCL2*, *CARD9*, and *NRPL3* suggests that ZIKV could lead to apoptosis and activation of inflammasomes in placental cells [66,77–79].

In parallel, at term placenta from CZS cases showed a decrease in *ISG15* mRNA, which was already identified as being protective of CZS ocular manifestations [80]. Another role of ISG15 is to modulate IFN responses since IFNλ4 blocks type I IFN response using the ISG15 and USP18 ubiquitin system [33]. Hence, we can hypothesize that a genetically regulated well-adjusted production of type I and type III IFN levels could lead to a proper protective response to ZIKV infection in the placenta, which could prevent CZS severe outcomes (Fig. 5). Nevertheless, either genetic or functional findings should be independently replicated in other populations.

**Fig 5.**
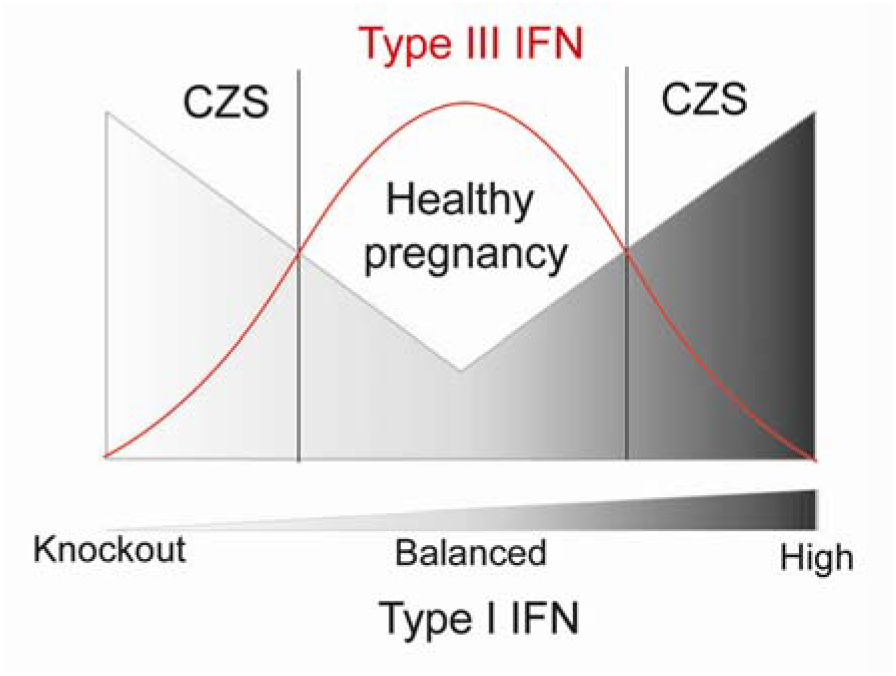
Schematic representation of the relation between Type I and III IFN in congenital ZIKV infections regarding abnormal CZS, showing that lower risk of developing CZS is related to higher levels of IFNL and balanced levels of Type-I IFN.

In summary, our study showed that intensity of immune responses during ZIKV infections in humans can be regulated by host genes. During pregnancy, genetic regulatory pathways control placental tissue-specific type I and type III IFN expression during ZIKV congenital infection influencing fetal neurological damage. Understanding of this novel pathway may help in the development of a custom pharmacological intervention to normalize its levels, which would likely affect and disrupt CNS development.

## Acknowledgements

The authors would like to thank Ms. Natália Pedra Gonçalves and Dr. Erica Louro da Fonseca from Vice Diretoria de Qualidade, Biomanguinhos, Fiocruz, for their help with ancestry analysis. We would also like to thank the team of Laboratório de Tecnologia Imunológica, Vice Diretoria de Desenvolvimento de Biomanguinhos, Fiocruz, for their technical and management support. This work was supported by Biomanguinhos and Instituto Oswaldo Cruz, Fiocruz, Brazil.

## Author Contributions

Conceptualization: MOM, ZV, PCCN, and TA; Data curation: TA, DPC, EAP, LCG, LG, MELM, and ZV; Formal analysis: TA, FK, AMVS, CCC, MRA, TLC, OCL, FSNM, LPF and EFC. Investigation: AMVS, CB, JS, AFS. Funding acquisition and Resources: MM and MOM. Supervision: PCCN, ZV and MOM; Writing – original draft: TA; Writing – review & editing: DPC, PCCN, ZV, and MOM.

## Competing interests

The authors declare no competing interests.

## Data availability

The data that support the findings of this study are available from the corresponding author upon reasonable request.

## Notes

### Competing Interest Statement

The authors have declared no competing interest.

### Summary of Updates

Dear readers, The authors have decided to revise their manuscript entitled Congenital Zika Syndrome is associated with maternal genetic background, in the pre-print platform. The authors have improved the genetic analysis by increasing the sample size, also investigating other target genes. After these updates, the genetic association as presented in the past preprint version was not confirmed, but a novel association was identified, and now it is a final contribution for these analyses. If you have any questions, please contact the corresponding author Dr. Milton Moraes. Best Regards, Milton Moraes

