## Supplementary table 1 for "Congenital Zika Syndrome is associated with interferon alfa receptor 1"

**Supplementary Table 1. Selected SNPs description**

|  |  |  |  |  | Allele frequencies (A1) | |  | Information from annotation | |  |
| --- | --- | --- | --- | --- | --- | --- | --- | --- | --- | --- |
| Chr | SNP ID | (GRCh37/hg19) | A1 | A2 | Africans | Europeans |  | Gene | Func.refGene | Reference |
| 21 | rs2257167 | 34715699 | G | C | 0.809 | 0.850 |  | *IFNAR1* | Missense Variant | ^31, 32^ |
| 21 | rs2843710 | 34696707 | C | G | 0.650 | 0.590 |  | *IFNAR1* | Upstream | ^31, 32^ |
| 19 | rs12979860 | 39738787 | C | T | 0.340 | 0.691 |  | *IFNL4* | Intronic | ^49^ |
| 19 | rs4803222 | 39739353 | C | G | 0.251 | 0.296 |  | *IFNL4* | 5’ UTR |  |
| 19 | rs8109886 | 39742762 | C | A | 0.148 | 0.567 |  | *IFNL4, IFNL2* | Intergenic | ^29^ |
| 19 | rs8099917 | 39743165 | G | T | 0.036 | 0.168 |  | *IFNL4, IFNL2* | Intergenic | ^49,65^ |

Annotation information from ANNOVAR with references for previous association with infectious disease.
