## Supplementary table 2 for "Congenital Zika Syndrome is associated with interferon alfa receptor 1"

**Supplementary Table 2. Selected SNP allele descriptions**

| *IFNAR1* | | | | | |
| --- | --- | --- | --- | --- | --- |
| rs2843710 | rs2257167 | rs17875834 | rs2834202 | Haplotype frequencies | |
|  |  |  |  | Africans | Europeans |
| C | G | C | A | 0.421 | 0.581 |
| C | G | T | A | 0.245 | 0.000 |
| G | C | C | A | 0.153 | 0.000 |
| G | G | C | G | 0.109 | 0.277 |
| C | G | C | G | 0.051 | 0.000 |
| G | G | C | A | 0.019 | 0.000 |
| G | C | C | A | 0.000 | 0.125 |
| *IFNL2-4* | | | | | |
| rs12979860 | rs4803222 | rs8109886 | rs8099917 | Haplotype frequencies | |
|  |  |  |  | Africans | Europeans |
| T | G | A | T | 0.406 | 0.017 |
| T | C | A | T | 0.215 | 0.127 |
| C | G | A | T | 0.195 | 0.121 |
| C | G | C | T | 0.145 | 0.566 |
| T | C | A | G | 0.036 | 0.165 |

Alelle frequencies among Africans and Europeans from the 1000 Genomes Project
