## Supplementary table 3 for "Congenital Zika Syndrome is associated with interferon alfa receptor 1"

**Supplementary Table 3. Association with *IFNAR1* haplotypes and CZS in newborns during ZIKV congenital infections**

|  |  |  |  | No adjustments |  | Adjusted by trimester of exposure to ZIKV and ancestrality | | |
| --- | --- | --- | --- | --- | --- | --- | --- | --- |
|  |  | No CZS findings | Abnormal CZS | OR; p-value | | |  |  |
| rs2257167 | rs2843710 | (%) | (%) | CI (95%) | | |  | FDR p-value |
| G | C | 62.30 | 53.72 | ref |  | ref |  | ref |
| C | C | 1.12 | 3.55 | 1.86; 0.5248 |  | 2.88; 0.3174 |  | 0.635 |
|  |  |  |  | (0.27-12.51) |  | (0.36-22.85) |  |  |
| C | G | 12.10 | 21.58 | 2.15; 0.0571 |  | 2.85; 0.021 |  | 0.084 |
|  |  |  |  | (0.98-4.70) |  | (1.18-6.88) |  |  |
| G | G | 24.48 | 21.16 | 0.88; 0.7150 |  | 0.97; 0.9491 |  | 1.000 |
|  |  |  |  | (0.45-1.73) |  | (0.45-2.12) |  |  |

DNA samples from 143 newborns with congenital infection by ZIKV, which developed CZS or not, were genotyped for rs2257167 and rs2843710 SNPs at *IFNAR1*. Data were adjusted considering the trimester of the first symptoms or asymptomatic ZIKV infections and the percentage of the African and European ancestry of each individual. The total number of the genotyped samples for each SNP may vary due to genotype miscalling. Haplotype G/C (rs2257167/rs2843710) was used as haplobase. Odds ratio (OR) with 95% confidence interval (CI) and p-values. We conducted Type I error adjustment of multiple comparisons by the False Discovery Rate (FDR) method.
