## Supplementary table 4 for "Congenital Zika Syndrome is associated with interferon alfa receptor 1"

**Supplementary Table 4. Association study with mother *IFNAR1* and *IFNL* SNPs and CZS abnormalities**

|  | No CZS findings | |  | Abnormal CZS | |  | Adjusted by trimester of ZIKV exposure and ancestrality | | | | |
| --- | --- | --- | --- | --- | --- | --- | --- | --- | --- | --- | --- |
|  | N^a^ | % |  | N^a^ | % |  | OR | lower | upper | p-value | FDR p-value |
| rs2257167 | | | | | | | | | | | |
| G/G | 50 | 76.9 |  | 43 | 69.4 |  | ref |  |  | 0.2879 | 0.431850 |
| C/G | 13 | 20 |  | 15 | 24.2 |  | 2.19 | 0.77 | 6.22 |  |  |
| C/C | 2 | 3.1 |  | 4 | 6.5 |  | 1.92 | 0.28 | 13.06 |  |  |
| G/G | 50 | 76.9 |  | 43 | 69.4 |  | ref |  |  | 0.1156 | 0.307320 |
| C/G-C/C | 15 | 23.1 |  | 19 | 30.6 |  | 2.13 | 0.82 | 5.56 |  |  |
| G/G-C/G | 63 | 96.9 |  | 58 | 93.5 |  | ref |  |  | 0.6105 | 0.732588 |
| C/C | 2 | 3.1 |  | 4 | 6.5 |  | 1.63 | 0.24 | 10.85 |  |  |
| G/G-C/C | 52 | 80 |  | 47 | 75.8 |  | ref |  |  | 0.1537 | 0.307320 |
| C/G | 13 | 20 |  | 15 | 24.2 |  | 2.10 | 0.75 | 5.92 |  |  |
| Additive | 65 | 51.2 |  | 62 | 48.8 |  | 1.72 | 0.80 | 3.67 | 0.1528 | 0.307320 |
| rs2843710 | | | | | | | | | | | |
| G/G | 26 | 40.6 |  | 27 | 43.5 |  | ref |  |  | 0.9454 | 1 |
| C/G | 26 | 40.6 |  | 26 | 41.9 |  | 1.06 | 0.43 | 2.60 |  |  |
| C/C | 12 | 18.8 |  | 9 | 14.5 |  | 0.87 | 0.27 | 2.77 |  |  |
| C/C | 26 | 40.6 |  | 27 | 43.5 |  | ref |  |  | 0.9964 | 1 |
| C/G-G/G | 38 | 59.4 |  | 35 | 56.5 |  | 1.00 | 0.44 | 2.29 |  |  |
| C/C-C/G | 52 | 81.2 |  | 53 | 85.5 |  | ref |  |  | 0.7587 | 1 |
| G/G | 12 | 18.8 |  | 9 | 14.5 |  | 0.85 | 0.29 | 2.46 |  |  |
| C/C-G/G | 38 | 59.4 |  | 36 | 58.1 |  | ref |  |  | 0.8079 | 1 |
| C/G | 26 | 40.6 |  | 26 | 41.9 |  | 1.11 | 0.49 | 2.53 |  |  |
| Additive | 64 | 50.8 |  | 62 | 49.2 |  | 0.96 | 0.55 | 1.67 | 0.8748 | 1 |
| rs12979860 | | | | | | | | | | | |
| C/C | 28 | 36.4 |  | 29 | 38.2 |  | ref |  |  | 0.23315 | 0.498380 |
| C/T | 35 | 45.5 |  | 40 | 52.6 |  | 1.05 | 0.47 | 2.34 |  |  |
| T/T | 14 | 18.2 |  | 7 | 9.2 |  | 0.39 | 0.12 | 1.35 |  |  |
| C/C | 28 | 36.4 |  | 29 | 38.2 |  | ref |  |  | 0.69046 | 0.828552 |
| C/T-T/T | 49 | 63.6 |  | 47 | 61.8 |  | 0.86 | 0.40 | 1.84 |  |  |
| C/C-C/T | 63 | 81.8 |  | 69 | 90.8 |  | ref |  |  | 0.08861 | 0.498380 |
| T/T | 14 | 18.2 |  | 7 | 9.2 |  | 0.38 | 0.12 | 1.20 |  |  |
| C/C-T/T | 42 | 54.5 |  | 36 | 47.4 |  | ref |  |  | 0.43751 | 0.656265 |
| C/T | 35 | 45.5 |  | 40 | 52.6 |  | 1.34 | 0.64 | 2.79 |  |  |
| Additive | 77 | 50.3 |  | 76 | 49.7 |  | 0.72 | 0.41 | 1.26 | 0.24919 | 0.498380 |
| rs8099917 | | | | | | | | | | | |
| T/T | 54 | 71.1 |  | 48 | 65.8 |  | ref |  |  | 0.5761 | 0.882585 |
| G/T | 17 | 22.4 |  | 22 | 30.1 |  | 1.49 | 0.62 | 3.57 |  |  |
| G/G | 5 | 6.6 |  | 3 | 4.1 |  | 0.72 | 0.14 | 3.63 |  |  |
| T/T | 54 | 71.1 |  | 48 | 65.8 |  | ref |  |  | 0.5280 | 0.882585 |
| G/T-G/G | 22 | 28.9 |  | 25 | 34.2 |  | 1.30 | 0.58 | 2.90 |  |  |
| T/T-G/T | 71 | 93.4 |  | 70 | 95.9 |  | ref |  |  | 0.5884 | 0.882585 |
| G/G | 5 | 6.6 |  | 3 | 4.1 |  | 0.65 | 0.13 | 3.19 |  |  |
| T/T-G/G | 59 | 77.6 |  | 51 | 69.9 |  | ref |  |  | 0.3308 | 0.882585 |
| G/T | 17 | 22.4 |  | 22 | 30.1 |  | 1.53 | 0.65 | 3.63 |  |  |
| Additive | 76 | 51 |  | 73 | 49 |  | 1.09 | 0.59 | 2.03 | 0.7858 | 0.942948 |
| rs8109886 | | | | | | | | | | | |
| A/A | 29 | 38.2 |  | 19 | 26.8 |  | ref |  |  | 0.3743 | 0.561420 |
| C/A | 38 | 50 |  | 37 | 52.1 |  | 1.47 | 0.62 | 3.46 |  |  |
| C/C | 9 | 11.8 |  | 15 | 21.1 |  | 2.25 | 0.70 | 7.20 |  |  |
| A/A | 29 | 38.2 |  | 19 | 26.8 |  | ref |  |  | 0.2445 | 0.549480 |
| C/A-C/C | 47 | 61.8 |  | 52 | 73.2 |  | 1.62 | 0.71 | 3.69 |  |  |
| A/A-C/A | 67 | 88.2 |  | 56 | 78.9 |  | ref |  |  | 0.2747 | 0.549480 |
| C/C | 9 | 11.8 |  | 15 | 21.1 |  | 1.76 | 0.63 | 4.92 |  |  |
| A/A-C/C | 38 | 50 |  | 34 | 47.9 |  | ref |  |  | 0.7900 | 0.947976 |
| C/A | 38 | 50 |  | 37 | 52.1 |  | 1.11 | 0.52 | 2.35 |  |  |
| Additive | 76 | 51.7 |  | 71 | 48.3 |  | 1.49 | 0.85 | 2.63 | 0.1612 | 0.549480 |
| rs4803222 | | | | | | | | | | | |
| G/G | 35 | 48.6 |  | 34 | 46.6 |  | ref |  |  | 0.6989 | 0.938808 |
| C/G | 29 | 40.3 |  | 33 | 45.2 |  | 1.39 | 0.62 | 3.09 |  |  |
| C/C | 8 | 11.1 |  | 6 | 8.2 |  | 0.97 | 0.25 | 3.82 |  |  |
| G/G | 35 | 48.6 |  | 34 | 46.4 |  | ref |  |  | 0.4965 | 0.938808 |
| C/G-C/C | 37 | 51.4 |  | 39 | 53.4 |  | 1.30 | 0.61 | 2.79 |  |  |
| G/G-C/G | 64 | 88.9 |  | 67 | 91.8 |  | ref |  |  | 0.7823 | 0.938808 |
| C/C | 8 | 11.1 |  | 6 | 8.2 |  | 0.83 | 0.22 | 3.09 |  |  |
| G/G-C/C | 43 | 59.7 |  | 40 | 54.8 |  | ref |  |  | 0.3979 | 0.938808 |
| C/G | 29 | 40.3 |  | 33 | 45.2 |  | 1.39 | 0.64 | 3.01 |  |  |
| Additive | 72 | 49.7 |  | 73 | 50.3 |  | 1.13 | 0.63 | 2.03 | 0.6887 | 0.938808 |

DNA samples from 153 mothers with congenital infection by ZIKV, with CZS or not, were genotyped for *IFNAR1* SNPs rs2257167 and rs2843710, and *IFNL* rs8109886, rs12979860, rs8099917 and rs4803222. Data were adjusted considering the trimester of the first symptoms or asymptomatic ZIKV infections and the percentage of the African and European ancestry of each individual. The total number of genotyped samples for each SNP may vary due to genotype miscalling. Major allele was used as baseline. Odds ratio (OR) with 95% confidence interval (CI) and p-values. We conducted Type I error adjustment of multiple comparisons by the False Discovery Rate (FDR) method.

parisons by the Holm-Bonferroni method. P-values ** ≤ 0.01, * ≤ 0.05, and ≤ 0.1.
