## Supplementary table 5 for "Congenital Zika Syndrome is associated with interferon alfa receptor 1"

**Supplementary Table 5. *IFNL2-4* SNP interaction in mother-child pairs and association with cephalic alteration outcomes**

|  | rs2257167 | rs2843710 | rs12979860 | rs8109886 | rs8099917 | rs4803222 |
| --- | --- | --- | --- | --- | --- | --- |
| logLikeRatio | 4.71 | 2.89 | 4.17 | 2.06 | 2.55 | 1.40 |
| p-values | 0.193 | 0.407 | 0.242 | 0.558 | 0.466 | 0.703 |

The DNA samples from 42 pairs of ZIKV-infected mothers and their respective newborns, with or without CZS were genotyped for IFNL2-4 SNPs rs8109886, rs12979860, rs8099917, and rs4803222. EMIM analysis using a multinomial model to test the existence of (and estimate) the genotype relative risk parameters that increase (or decrease) the possibility that a child will be affected. No significant effects of mother or child nor mother-child genotype interaction in cephalic alteration outcomes were observed using this sample size.
