## Supplementary table 6 for "Congenital Zika Syndrome is associated with interferon alfa receptor 1"

**Supplementary Table 6. Demographic and clinical features of pregnant woman enrolled in placental gene expression analyses**

|  |  | Congenital ZIKV infection (n=74) | | |  | Healthy pregnancy (n=10) |
| --- | --- | --- | --- | --- | --- | --- |
|  |  | No CZS findings (n=45) |  | Abnormal CZS (n=29) |  |  |
| Mothers’ age | < 40 | 33 (73.33%) |  | 26(89.66%) |  | 10 (100%) |
|  | ≥40 | 11(24.44%) |  | 0(0%) |  | 0 (0%) |
|  | Unknow | 1(2.22%) |  | 3(10.34%) |  | 0 (0%) |
| Trimester of exposure to ZIKV | 1st | 12(26.67%) |  | 18(62.07%) |  | NA |
|  | 2nd | 20(44.44%) |  | 1(3.45%) |  | NA |
|  | 3rd | 9(20.00%) |  | 0(0%) |  | NA |
|  | Asymptomatic | 1(2.22%) |  | 5(17.24%) |  | NA |
|  | Unknow | 3(6.67%) |  | 5(17.24%) |  | NA |
| ZIKV PCR in placenta | Positive | 11(24.44%) |  | 9(31.03%) |  | 0 (0%) |
|  | Negative | 34(75.56%) |  | 20(68.97%) |  | 10 (100%) |
