## Supplementary table 7 for "Congenital Zika Syndrome is associated with interferon alfa receptor 1"

**Supplementary Table 7. Placental gene expression according to demographic and clinical features**

|  | Mothers’ age | |  | ZIKV PCR in placenta | |  | Trimester of ZIKV infection | | | |
| --- | --- | --- | --- | --- | --- | --- | --- | --- | --- | --- |
|  | < 40 | ≥40 |  | Negative | Positive |  | 1^st^ trimester | 2nd trimester | 3nd trimestre | Asymptomatic |
| *AIM2* | 0.42 ± 0.15 | 0.28 ± 0.11 |  | 0.35 ± 0.13 | 0.52 ± 0.12 |  | 0.39 ± 0.16 | 0.39 ± 0.17 | 0.41 ± 0.16 | 0.38 ± 0.09 |
| *BCL2* | 0.21 ± 0.07 | 0.19 ± 0.04 |  | 0.20 ± 0.06 | 0.23 ± 0.07 |  | 0.21 ± 0.09 | 0.20 ± 0.04 | 0.21 ± 0.04 | 0.25 ± 0.11 |
| *CARD6* | 0.28 ± 0.13 | 0.23 ± 0.05 |  | 0.25 ± 0.13 | 0.29 ± 0.10 |  | 0.24 ± 0.09 | 0.26 ± 0.09 | 0.32 ± 0.09 | 0.36 ± 0.33 |
| *CARD9* | 0.47 ± 0.23 | 0.42 ± 0.23 |  | 0.43 ± 0.23 | 0.56 ± 0.20 |  | 0.44 ± 0.22 | 0.49 ± 0.19 | 0.48 ± 0.16 | 0.60 ± 0.48 |
| *CCL22* | 0.59 ± 0.39 | 0.35 ± 0.00 |  | 0.60 ± 0.40 | 0.54 ± 0.26 |  | 0.67 ± 0.34 | 0.46 ± 0.23 | 0.65 ± 0.36 | 0.73 ± 0.81 |
| *CCR2* | 0.22 ± 0.08 | 0.20 ± 0.03 |  | 0.21 ± 0.09 | 0.23 ± 0.03 |  | 0.21 ± 0.08 | 0.21 ± 0.03 | 0.22 ± 0.04 | 0.30 ± 0.19 |
| *CCR3* | 0.22 ± 0.05 | 0.20 ± 0.02 |  | 0.20 ± 0.05 | 0.24 ± 0.03 |  | 0.21 ± 0.03 | 0.21 ± 0.03 | 0.23 ± 0.05 | 0.28 ± 0.12 |
| *CCR4* | 0.26 ± 0.10 | 0.21 ± 0.02 |  | 0.24 ± 0.10 | 0.27 ± 0.05 |  | 0.23 ± 0.05 | 0.25 ± 0.07 | 0.28 ± 0.07 | 0.37 ± 0.25 |
| *CCR5* | 0.25 ± 0.11 | 0.23 ± 0.04 |  | 0.24 ± 0.11 | 0.26 ± 0.04 |  | 0.24 ± 0.10 | 0.24 ± 0.04 | 0.26 ± 0.05 | 0.38 ± 0.30 |
| *CD36* | 0.31 ± 0.15 | 0.27 ± 0.07 |  | 0.31 ± 0.13 | 0.31 ± 0.17 |  | 0.31 ± 0.17 | 0.27 ± 0.10 | 0.33 ± 0.07 | 0.40 ± 0.28 |
| *CLEC5A* | 0.30 ± 0.13 | 0.26 ± 0.07 |  | 0.28 ± 0.11 | 0.32 ± 0.13 |  | 0.30 ± 0.14 | 0.27 ± 0.06 | 0.29 ± 0.04 | 0.39 ± 0.26 |
| *DAP12* | 0.36 ± 0.18 | 0.32 ± 0.10 |  | 0.35 ± 0.15 | 0.37 ± 0.20 |  | 0.36 ± 0.16 | 0.33 ± 0.12 | 0.36 ± 0.12 | 0.48 ± 0.46 |
| *DCSIGN* | 0.31 ± 0.13 | 0.27 ± 0.07 |  | 0.29 ± 0.11 | 0.34 ± 0.12 |  | 0.31 ± 0.14 | 0.29 ± 0.07 | 0.31 ± 0.06 | 0.39 ± 0.24 |
| *FOXP3* | 0.31 ± 0.18 | 0.23 ± 0.03 |  | 0.29 ± 0.19 | 0.32 ± 0.06 |  | 0.28 ± 0.09 | 0.26 ± 0.08 | 0.27 ± 0.04 | 0.60 ± 0.56 |
| *IDO* | 0.42 ± 0.26 | 0.21 ± 0.06 |  | 0.36 ± 0.25 | 0.45 ± 0.23 |  | 0.43 ± 0.28 | 0.32 ± 0.16 | 0.39 ± 0.16 | 0.45 ± 0.49 |
| *IDO2* | 0.30 ± 0.16 | 0.21 ± 0.05 |  | 0.26 ± 0.14 | 0.33 ± 0.17 |  | 0.28 ± 0.16 | 0.27 ± 0.10 | 0.29 ± 0.11 | 0.40 ± 0.34 |
| *IFI16* | 0.40 ± 0.20 | 0.34 ± 0.10 |  | 0.38 ± 0.17 | 0.42 ± 0.21 |  | 0.39 ± 0.20 | 0.37 ± 0.13 | 0.43 ± 0.13 | 0.51 ± 0.41 |
| *IFI27* | 0.32 ± 0.18 | 0.21 ± 0.06 |  | 0.28 ± 0.16 | 0.35 ± 0.17 |  | 0.31 ± 0.16 | 0.27 ± 0.12 | 0.32 ± 0.12 | 0.43 ± 0.39 |
| *IFI35* | 0.35 ± 0.22 | 0.28 ± 0.08 |  | 0.34 ± 0.20 | 0.35 ± 0.21 |  | 0.35 ± 0.19 | 0.31 ± 0.15 | 0.33 ± 0.07 | 0.54 ± 0.56 |
| *IFI44* | 0.42 ± 0.23 | 0.31 ± 0.07 |  | 0.38 ± 0.19 | 0.45 ± 0.24 |  | 0.40 ± 0.20 | 0.37 ± 0.15 | 0.45 ± 0.15 | 0.56 ± 0.54 |
| *IFI6* | 0.31 ± 0.11 | 0.25 ± 0.07 |  | 0.30 ± 0.10 | 0.32 ± 0.13 |  | 0.30 ± 0.11 | 0.28 ± 0.10 | 0.31 ± 0.11 | 0.36 ± 0.22 |
| *IFIH1* | 0.40 ± 0.23 | 0.30 ± 0.07 |  | 0.37 ± 0.20 | 0.40 ± 0.24 |  | 0.39 ± 0.22 | 0.33 ± 0.12 | 0.42 ± 0.12 | 0.52 ± 0.54 |
| *IFIT2* | 0.66 ± 0.42 | 0.46 ± 0.00 |  | 0.62 ± 0.40 | 0.63 ± 0.12 |  | 0.55 ± 0.17 | 0.46 ± 0.00 | 0.00 ± 0.00 | 1.14 ± 0.96 |
| *IFIT5* | 0.18 ± 0.04 | 0.18 ± 0.02 |  | 0.18 ± 0.04 | 0.19 ± 0.03 |  | 0.18 ± 0.05 | 0.18 ± 0.03 | 0.19 ± 0.02 | 0.23 ± 0.07 |
| *IFNL1* | 0.65 ± 0.39 | 0.85 ± 0.43 |  | 0.59 ± 0.33 | 0.85 ± 0.46 |  | 0.57 ± 0.40 | 0.71 ± 0.36 | 0.99 ± 0.45 | 0.67 ± 0.24 |
| *IFNL2* | 1.28 ± 0.83 | 1.53 ± 0.68 |  | 1.17 ± 0.67 | 1.69 ± 0.95 |  | 1.18 ± 0.90 | 1.17 ± 0.64 | 1.83 ± 0.62 | 1.82 ± 1.06 |
| *IFNL3* | 0.44 ± 0.22 | 0.57 ± 0.27 |  | 0.42 ± 0.19 | 0.55 ± 0.24 |  | 0.38 ± 0.15 | 0.49 ± 0.27 | 0.67 ± 0.24 | 0.48 ± 0.08 |
| *IFNL4* | 0.54 ± 0.29 | 0.70 ± 0.36 |  | 0.50 ± 0.27 | 0.69 ± 0.29 |  | 0.43 ± 0.20 | 0.62 ± 0.33 | 0.85 ± 0.33 | 0.62 ± 0.21 |
| *IFNA1* | 0.25 ± 0.08 | 0.23 ± 0.04 |  | 0.24 ± 0.08 | 0.27 ± 0.03 |  | 0.24 ± 0.07 | 0.25 ± 0.05 | 0.25 ± 0.04 | 0.33 ± 0.18 |
| *IFNAR* | 0.39 ± 0.21 | 0.27 ± 0.08 |  | 0.36 ± 0.18 | 0.39 ± 0.23 |  | 0.38 ± 0.22 | 0.34 ± 0.12 | 0.40 ± 0.13 | 0.46 ± 0.44 |
| *IFNB* | 0.36 ± 0.10 | 0.35 ± 0.14 |  | 0.35 ± 0.10 | 0.41 ± 0.10 |  | 0.35 ± 0.10 | 0.37 ± 0.13 | 0.40 ± 0.09 | 0.35 ± 0.08 |
| *IFNG* | 0.46 ± 0.25 | 0.31 ± 0.07 |  | 0.42 ± 0.24 | 0.48 ± 0.18 |  | 0.42 ± 0.12 | 0.39 ± 0.14 | 0.50 ± 0.17 | 0.60 ± 0.64 |
| *IFTM1* | 0.31 ± 0.11 | 0.28 ± 0.06 |  | 0.30 ± 0.10 | 0.30 ± 0.13 |  | 0.31 ± 0.12 | 0.28 ± 0.08 | 0.34 ± 0.08 | 0.35 ± 0.18 |
| *IFTM3* | 0.20 ± 0.04 | 0.20 ± 0.01 |  | 0.20 ± 0.04 | 0.20 ± 0.01 |  | 0.20 ± 0.02 | 0.20 ± 0.01 | 0.21 ± 0.01 | 0.25 ± 0.12 |
| *IL10* | 0.33 ± 0.12 | 0.31 ± 0.10 |  | 0.32 ± 0.12 | 0.36 ± 0.08 |  | 0.31 ± 0.08 | 0.32 ± 0.10 | 0.35 ± 0.07 | 0.44 ± 0.29 |
| *IL12* | 0.72 ± 0.37 | 0.46 ± 0.00 |  | 0.67 ± 0.36 | 0.74 ± 0.33 |  | 0.77 ± 0.37 | 0.53 ± 0.15 | 0.61 ± 0.20 | 0.91 ± 0.68 |
| *IL17* | 0.46 ± 0.38 | 0.48 ± 0.24 |  | 0.47 ± 0.36 | 0.48 ± 0.25 |  | 0.34 ± 0.14 | 0.40 ± 0.28 | 0.55 ± 0.31 | 1.53 ± 0.00 |
| *IL18* | 0.30 ± 0.11 | 0.28 ± 0.09 |  | 0.29 ± 0.12 | 0.32 ± 0.07 |  | 0.29 ± 0.10 | 0.30 ± 0.09 | 0.31 ± 0.07 | 0.40 ± 0.27 |
| *IL1A* | 0.46 ± 0.26 | 0.16 ± 0.00 |  | 0.46 ± 0.30 | 0.38 ± 0.11 |  | 0.40 ± 0.07 | 0.32 ± 0.18 | 0.53 ± 0.00 | 0.58 ± 0.52 |
| *IL1B* | 0.57 ± 0.25 | 0.20 ± 0.00 |  | 0.61 ± 0.26 | 0.48 ± 0.20 |  | 0.58 ± 0.10 | 0.41 ± 0.25 | 0.74 ± 0.00 | 0.62 ± 0.51 |
| *IL2* | 0.43 ± 0.16 | 0.32 ± 0.09 |  | 0.36 ± 0.10 | 0.51 ± 0.19 |  | 0.40 ± 0.19 | 0.43 ± 0.13 | 0.39 ± 0.13 | 0.33 ± 0.10 |
| *IL21* | 0.45 ± 0.21 | 0.00 ± 0.00 |  | 0.44 ± 0.22 | 0.63 ± 0.06 |  | 0.48 ± 0.22 | 0.48 ± 0.37 | 0.00 ± 0.00 | 0.00 ± 0.00 |
| *IL22RA* | 0.27 ± 0.11 | 0.25 ± 0.06 |  | 0.27 ± 0.11 | 0.29 ± 0.05 |  | 0.25 ± 0.05 | 0.27 ± 0.07 | 0.28 ± 0.05 | 0.40 ± 0.32 |
| *IL23A* | 0.39 ± 0.19 | 0.31 ± 0.12 |  | 0.35 ± 0.16 | 0.45 ± 0.22 |  | 0.36 ± 0.17 | 0.39 ± 0.22 | 0.48 ± 0.20 | 0.21 ± 0.03 |
| *IL23R* | 0.53 ± 0.28 | 0.00 ± 0.00 |  | 0.56 ± 0.34 | 0.47 ± 0.17 |  | 0.46 ± 0.20 | 0.48 ± 0.16 | 0.61 ± 0.03 | 0.66 ± 0.66 |
| *IL2RA* | 0.36 ± 0.22 | 0.19 ± 0.07 |  | 0.32 ± 0.21 | 0.39 ± 0.19 |  | 0.30 ± 0.11 | 0.29 ± 0.14 | 0.51 ± 0.22 | 0.53 ± 0.52 |
| *IL4* | 0.37 ± 0.19 | 0.17 ± 0.00 |  | 0.36 ± 0.21 | 0.32 ± 0.09 |  | 0.32 ± 0.05 | 0.26 ± 0.09 | 0.41 ± 0.00 | 0.46 ± 0.39 |
| *IL6* | 0.37 ± 0.17 | 0.17 ± 0.07 |  | 0.34 ± 0.18 | 0.36 ± 0.15 |  | 0.35 ± 0.12 | 0.28 ± 0.13 | 0.36 ± 0.14 | 0.44 ± 0.42 |
| *IL7* | 0.54 ± 0.25 | 0.30 ± 0.04 |  | 0.45 ± 0.26 | 0.51 ± 0.15 |  | 0.44 ± 0.17 | 0.41 ± 0.14 | 0.51 ± 0.19 | 0.87 ± 0.59 |
| *IL8* | 0.31 ± 0.18 | 0.22 ± 0.06 |  | 0.28 ± 0.17 | 0.31 ± 0.15 |  | 0.29 ± 0.16 | 0.25 ± 0.12 | 0.37 ± 0.12 | 0.38 ± 0.43 |
| *IP10* | 0.29 ± 0.12 | 0.26 ± 0.06 |  | 0.28 ± 0.11 | 0.32 ± 0.10 |  | 0.29 ± 0.11 | 0.28 ± 0.07 | 0.29 ± 0.05 | 0.38 ± 0.26 |
| *IRF3* | 0.48 ± 0.26 | 0.29 ± 0.10 |  | 0.45 ± 0.25 | 0.47 ± 0.24 |  | 0.48 ± 0.22 | 0.38 ± 0.17 | 0.50 ± 0.23 | 0.55 ± 0.59 |
| *IRF7* | 0.29 ± 0.09 | 0.27 ± 0.08 |  | 0.27 ± 0.08 | 0.32 ± 0.09 |  | 0.29 ± 0.10 | 0.28 ± 0.07 | 0.30 ± 0.05 | 0.34 ± 0.16 |
| *IRF9* | 0.26 ± 0.09 | 0.22 ± 0.04 |  | 0.24 ± 0.09 | 0.27 ± 0.07 |  | 0.25 ± 0.08 | 0.24 ± 0.04 | 0.25 ± 0.04 | 0.34 ± 0.22 |
| *ISG15* | 0.38 ± 0.27 | 0.39 ± 0.23 |  | 0.37 ± 0.25 | 0.40 ± 0.29 |  | 0.37 ± 0.24 | 0.36 ± 0.22 | 0.40 ± 0.14 | 0.64 ± 0.74 |
| *MARCO* | 0.44 ± 0.28 | 0.22 ± 0.03 |  | 0.36 ± 0.26 | 0.50 ± 0.24 |  | 0.40 ± 0.24 | 0.31 ± 0.16 | 0.38 ± 0.18 | 1.43 ± 0.00 |
| *MCP1* | 0.43 ± 0.22 | 0.29 ± 0.10 |  | 0.38 ± 0.18 | 0.47 ± 0.26 |  | 0.38 ± 0.20 | 0.41 ± 0.17 | 0.44 ± 0.17 | 0.57 ± 0.47 |
| *MIP1A* | 0.49 ± 0.27 | 0.33 ± 0.05 |  | 0.43 ± 0.23 | 0.55 ± 0.31 |  | 0.47 ± 0.31 | 0.42 ± 0.14 | 0.40 ± 0.08 | 0.63 ± 0.41 |
| *MIP1B* | 0.28 ± 0.13 | 0.23 ± 0.02 |  | 0.27 ± 0.13 | 0.30 ± 0.09 |  | 0.25 ± 0.07 | 0.25 ± 0.08 | 0.33 ± 0.09 | 0.45 ± 0.35 |
| *MMP2* | 0.27 ± 0.10 | 0.24 ± 0.04 |  | 0.25 ± 0.08 | 0.29 ± 0.11 |  | 0.26 ± 0.09 | 0.25 ± 0.05 | 0.28 ± 0.04 | 0.35 ± 0.22 |
| *MMP9* | 0.29 ± 0.16 | 0.19 ± 0.07 |  | 0.27 ± 0.15 | 0.28 ± 0.17 |  | 0.27 ± 0.13 | 0.24 ± 0.10 | 0.33 ± 0.10 | 0.36 ± 0.42 |
| *NOD2* | 0.28 ± 0.15 | 0.17 ± 0.06 |  | 0.25 ± 0.12 | 0.29 ± 0.17 |  | 0.27 ± 0.15 | 0.22 ± 0.08 | 0.28 ± 0.12 | 0.35 ± 0.30 |
| *NOS* | 0.31 ± 0.09 | 0.00 ± 0.00 |  | 0.30 ± 0.08 | 0.33 ± 0.08 |  | 0.36 ± 0.04 | 0.28 ± 0.12 | 0.41 ± 0.00 | 0.25 ± 0.06 |
| *NRLP1* | 0.34 ± 0.18 | 0.23 ± 0.02 |  | 0.31 ± 0.17 | 0.33 ± 0.16 |  | 0.37 ± 0.22 | 0.27 ± 0.06 | 0.32 ± 0.07 | 0.41 ± 0.35 |
| *NRLP3* | 0.43 ± 0.24 | 0.30 ± 0.03 |  | 0.35 ± 0.17 | 0.52 ± 0.28 |  | 0.42 ± 0.25 | 0.40 ± 0.16 | 0.45 ± 0.20 | 0.39 ± 0.38 |
| *OAS1* | 0.42 ± 0.18 | 0.38 ± 0.16 |  | 0.39 ± 0.16 | 0.45 ± 0.18 |  | 0.39 ± 0.17 | 0.41 ± 0.15 | 0.43 ± 0.09 | 0.49 ± 0.43 |
| *OAS2* | 0.37 ± 0.17 | 0.34 ± 0.16 |  | 0.36 ± 0.15 | 0.38 ± 0.19 |  | 0.38 ± 0.18 | 0.34 ± 0.15 | 0.51 ± 0.14 | 0.21 ± 0.13 |
| *OAS3* | 0.35 ± 0.15 | 0.28 ± 0.04 |  | 0.32 ± 0.13 | 0.40 ± 0.16 |  | 0.33 ± 0.13 | 0.33 ± 0.11 | 0.39 ± 0.11 | 0.47 ± 0.37 |
| *OASL* | 0.46 ± 0.25 | 0.37 ± 0.18 |  | 0.42 ± 0.23 | 0.51 ± 0.23 |  | 0.44 ± 0.23 | 0.43 ± 0.20 | 0.51 ± 0.19 | 0.58 ± 0.49 |
| *RANTES* | 0.34 ± 0.18 | 0.22 ± 0.04 |  | 0.36 ± 0.20 | 0.29 ± 0.09 |  | 0.32 ± 0.03 | 0.25 ± 0.08 | 0.35 ± 0.10 | 0.47 ± 0.40 |
| *RIG1* | 0.33 ± 0.14 | 0.27 ± 0.06 |  | 0.31 ± 0.11 | 0.34 ± 0.18 |  | 0.33 ± 0.16 | 0.31 ± 0.12 | 0.31 ± 0.07 | 0.38 ± 0.25 |
| *RIPK2* | 0.37 ± 0.19 | 0.26 ± 0.06 |  | 0.34 ± 0.16 | 0.38 ± 0.20 |  | 0.36 ± 0.20 | 0.33 ± 0.12 | 0.37 ± 0.11 | 0.44 ± 0.36 |
| *RNASEL* | 0.39 ± 0.24 | 0.26 ± 0.08 |  | 0.34 ± 0.19 | 0.43 ± 0.28 |  | 0.37 ± 0.24 | 0.33 ± 0.14 | 0.40 ± 0.16 | 0.50 ± 0.51 |
| *SOCS3* | 0.47 ± 0.23 | 0.41 ± 0.17 |  | 0.46 ± 0.23 | 0.44 ± 0.18 |  | 0.44 ± 0.22 | 0.46 ± 0.16 | 0.45 ± 0.15 | 0.65 ± 0.54 |
| *SOD2* | 0.36 ± 0.19 | 0.28 ± 0.07 |  | 0.34 ± 0.16 | 0.34 ± 0.21 |  | 0.36 ± 0.19 | 0.30 ± 0.13 | 0.35 ± 0.10 | 0.45 ± 0.40 |
| *STAT2* | 0.39 ± 0.22 | 0.36 ± 0.20 |  | 0.37 ± 0.20 | 0.41 ± 0.25 |  | 0.37 ± 0.20 | 0.36 ± 0.20 | 0.40 ± 0.14 | 0.55 ± 0.52 |
| *STING* | 0.45 ± 0.24 | 0.37 ± 0.17 |  | 0.42 ± 0.23 | 0.47 ± 0.21 |  | 0.43 ± 0.25 | 0.43 ± 0.19 | 0.45 ± 0.14 | 0.59 ± 0.52 |
| *TGFB* | 0.29 ± 0.09 | 0.25 ± 0.06 |  | 0.28 ± 0.09 | 0.30 ± 0.07 |  | 0.28 ± 0.08 | 0.28 ± 0.07 | 0.31 ± 0.08 | 0.36 ± 0.23 |
| *TICAM1* | 0.36 ± 0.26 | 0.27 ± 0.09 |  | 0.35 ± 0.25 | 0.34 ± 0.19 |  | 0.33 ± 0.19 | 0.28 ± 0.14 | 0.52 ± 0.18 | 0.56 ± 0.61 |
| *TLR2* | 0.44 ± 0.23 | 0.28 ± 0.10 |  | 0.38 ± 0.20 | 0.48 ± 0.26 |  | 0.40 ± 0.20 | 0.39 ± 0.15 | 0.45 ± 0.17 | 0.57 ± 0.56 |
| *TLR3* | 0.26 ± 0.08 | 0.24 ± 0.04 |  | 0.26 ± 0.09 | 0.26 ± 0.03 |  | 0.26 ± 0.07 | 0.25 ± 0.04 | 0.25 ± 0.03 | 0.34 ± 0.21 |
| *TLR4* | 0.41 ± 0.24 | 0.31 ± 0.07 |  | 0.38 ± 0.19 | 0.44 ± 0.27 |  | 0.39 ± 0.22 | 0.35 ± 0.15 | 0.40 ± 0.12 | 0.59 ± 0.55 |
| *TLR7* | 0.39 ± 0.27 | 0.32 ± 0.00 |  | 0.38 ± 0.27 | 0.40 ± 0.22 |  | 0.34 ± 0.14 | 0.28 ± 0.13 | 0.64 ± 0.11 | 0.70 ± 0.73 |
| *TLR8* | 0.45 ± 0.24 | 0.30 ± 0.05 |  | 0.39 ± 0.22 | 0.52 ± 0.19 |  | 0.40 ± 0.18 | 0.41 ± 0.14 | 0.49 ± 0.20 | 0.60 ± 0.72 |
| *TNFA* | 0.33 ± 0.19 | 0.20 ± 0.02 |  | 0.28 ± 0.18 | 0.35 ± 0.18 |  | 0.31 ± 0.19 | 0.27 ± 0.09 | 0.32 ± 0.12 | 0.36 ± 0.44 |
| *TNFRSF18* | 0.61 ± 0.34 | 0.35 ± 0.00 |  | 0.64 ± 0.42 | 0.56 ± 0.17 |  | 0.60 ± 0.14 | 0.48 ± 0.25 | 0.60 ± 0.18 | 0.80 ± 0.73 |
| *TNFSF15* | 0.26 ± 0.13 | 0.21 ± 0.08 |  | 0.24 ± 0.11 | 0.28 ± 0.14 |  | 0.25 ± 0.12 | 0.23 ± 0.08 | 0.26 ± 0.07 | 0.35 ± 0.28 |
| *TNFSF9* | 0.57 ± 0.30 | 0.29 ± 0.00 |  | 0.53 ± 0.34 | 0.57 ± 0.15 |  | 0.43 ± 0.15 | 0.53 ± 0.21 | 0.66 ± 0.16 | 1.09 ± 0.80 |
| *TYRO3* | 0.05 ± 0.07 | 0.03 ± 0.04 |  | 0.04 ± 0.07 | 0.04 ± 0.07 |  | 0.04 ± 0.06 | 0.05 ± 0.06 | 0.05 ± 0.10 | 0.07 ± 0.09 |

Gene expression values expressed in arbitrary units normalized by housekeeping genes selected by geNorm and NormFinder as well as *18S* ribosomal RNA and *RLP13* ribosomal protein L13.
