## Supplementary figure 1 for "Congenital Zika Syndrome is associated with interferon alfa receptor 1"

**A**

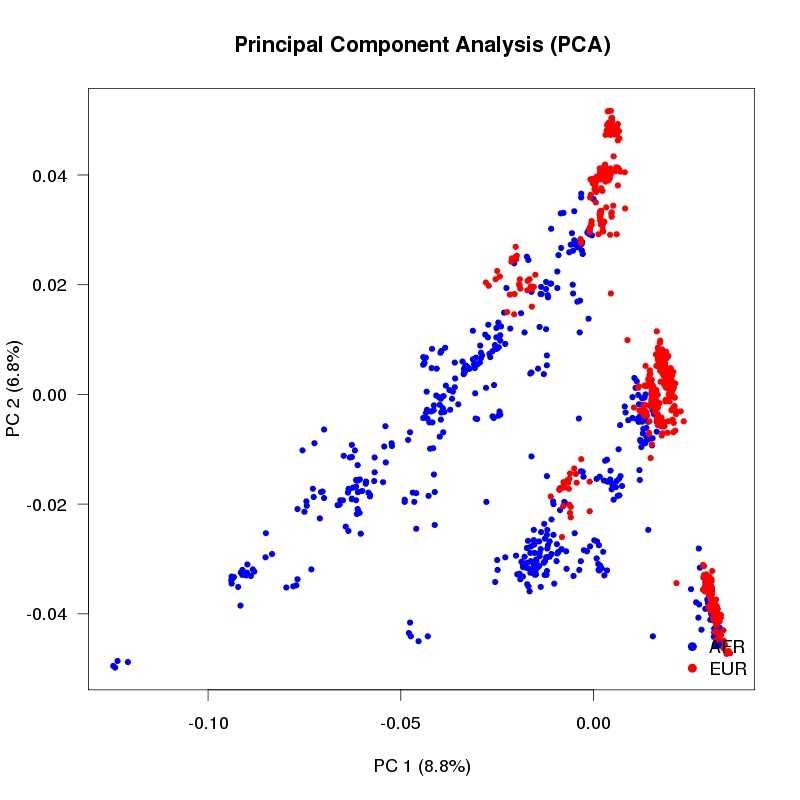

**B**

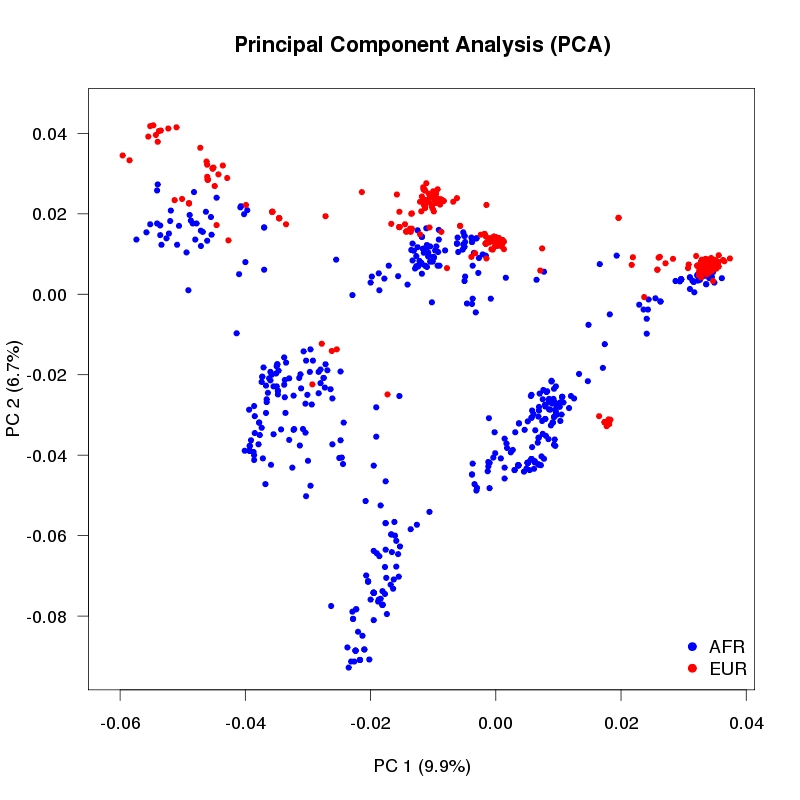

**C**

Africans	Europeans

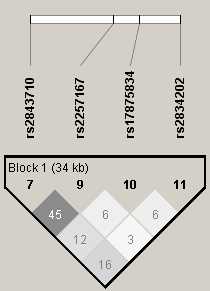

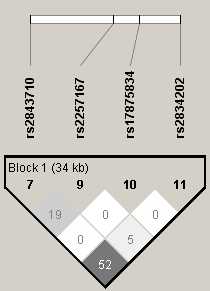

**D**

Africans	Europeans

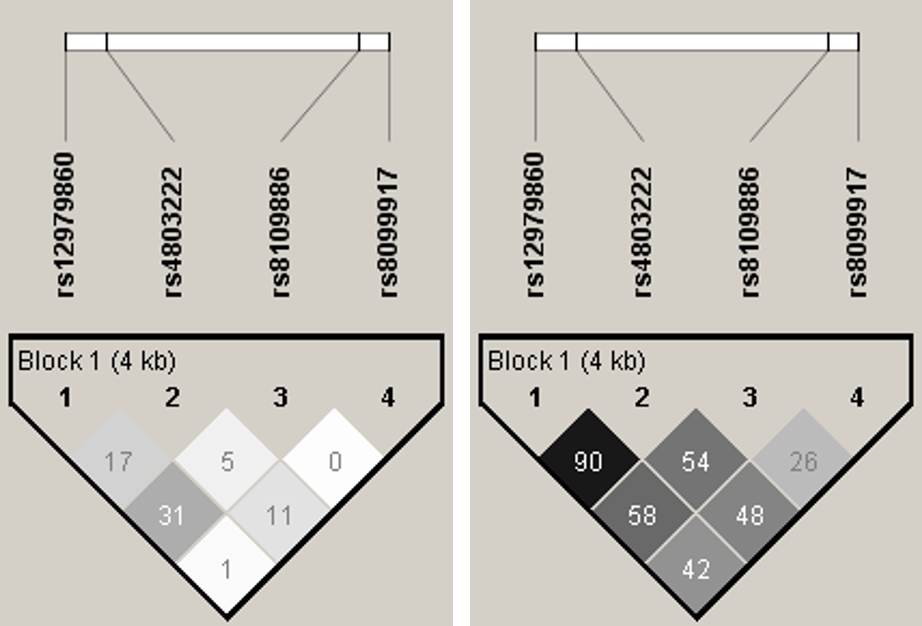

**Supplementary Fig. 1.** SNP selection and linkage disequilibrium analysis. **A and B**. Principal Component Analysis (PCA) of all SNPs located in the *IFNAR1* (A) and *IFNL* (B) regions which are found among African and European populations from the 1000 Genomes Project. PC-Principal Component. **C and D.** Pattern of linkage disequilibrium (LD) for the *IFNAR1* (C) and *IFNL2-4* (D) gene regions using data from African and European populations from phase 3 of the 1000 Genomes Project. Pairwise r2 values are shown in the diamonds.
