## Supplementary figure 2 for "Congenital Zika Syndrome is associated with interferon alfa receptor 1"

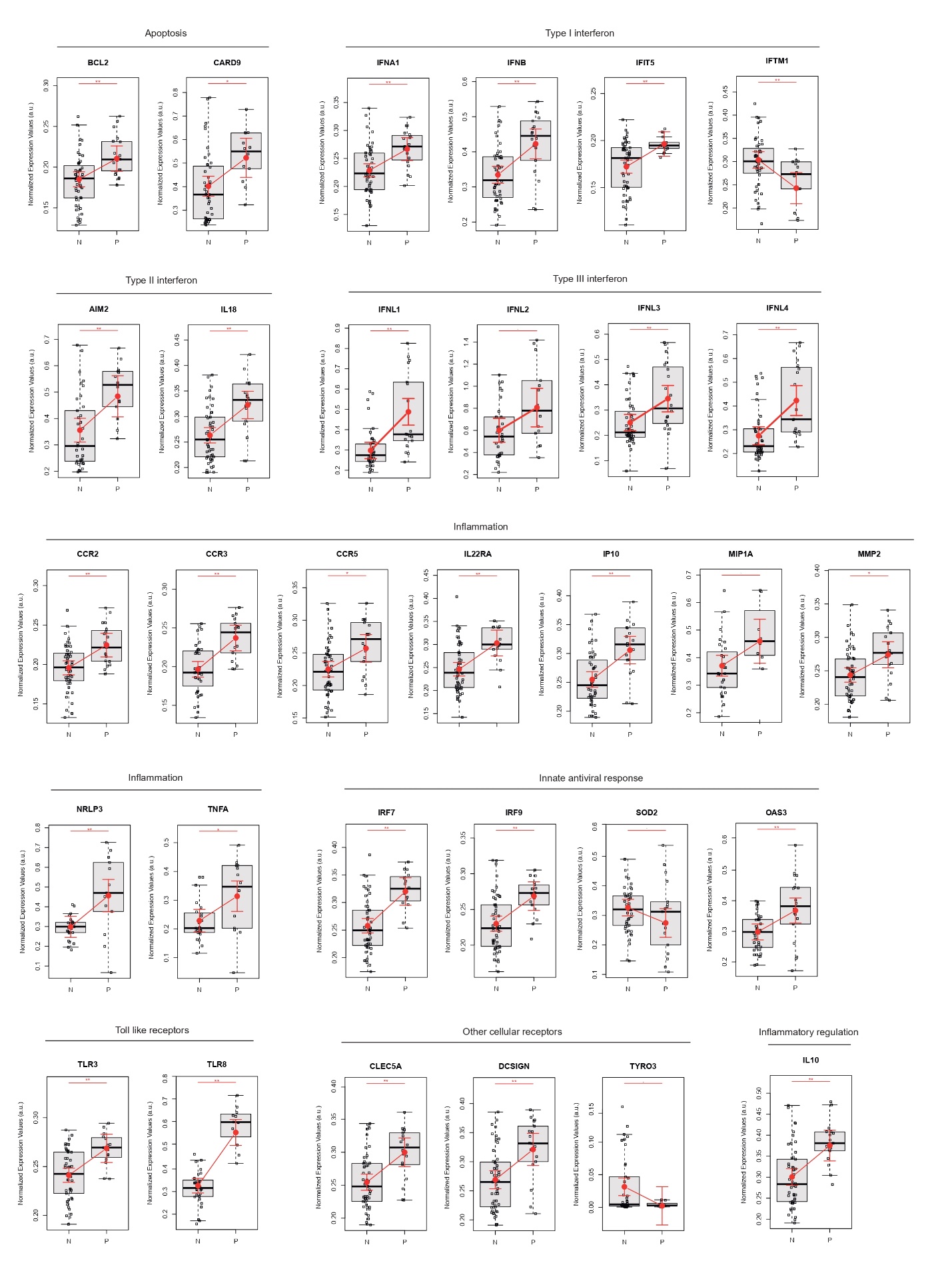

**Supplementary Fig. 2.** Gene expression profile of placenta negative and positive ZIKV PCR. Detailed graphs of differentially expressed genes in placenta negative (N; N=54) and positive (P; N=20) ZIKV PCR. Each dot corresponds to one placenta analyzed. The number of dots varies by gene analyzed due to failed amplifications. Median and standard deviation of gene expression values are normalized by housekeeping genes selected by the geNorm and NormFinder, as well as *18S* ribosomal RNA and*RLP13* ribosomal protein L13 (grey boxes). Values are adjusted by mothers’ age (below or equal to/above 40 years of age) and infection trimester (the trimester of pregnancy in which first Zika symptoms occur or asymptomatic ZIKV infections) (red lines). Pairwise comparisons of log-transformed (base 2) normalized expression means between/among the groups of interest were performed by contrasts/differences (fold-changes), obtained after both bi- and multivariate linear models were modified by ordinary least square regressions. Whenever the variable of interest had more than two levels, p-values were corrected by the Tukey Honest Significant Difference post-Hoc method. After gene-per-gene pairwise comparisons were carried out, we conducted a Type I error adjustment for the multiple comparisons by the Holm-Bonferroni method. P-values ** ≤ 0.01, * ≤ 0.05, and ≤ 0.1.
